# The atypical RNA-binding protein TAF15 regulates dorsoanterior neural development through diverse mechanisms in *Xenopus tropicalis*

**DOI:** 10.1101/2021.06.14.041913

**Authors:** Caitlin S. DeJong, Darwin S. Dichmann, Cameron R. T. Exner, Yuxiao Xu, Richard M. Harland

## Abstract

The FET family of atypical RNA-binding proteins includes Fused in sarcoma (Fus), Ewing’s sarcoma (EWS), and the TATA-binding protein-associate factor 15 (TAF15). FET proteins are highly conserved, suggesting specialized requirements for each protein. Fus regulates splicing of transcripts required for mesoderm differentiation and cell adhesion in *Xenopus*, but roles that EWS and TAF15 play remain unknown. Here we analyze the roles of maternally deposited and zygotically transcribed TAF15, which is essential for the proper development of dorsoanterior neural tissues. By measuring changes in exon usage and transcript abundance from TAF15-depleted embryos we found TAF15 may regulate dorsoanterior neural development through *fgfr4* and *ventx2*.*1*. TAF15 uses distinct mechanisms to downregulate FGFR4 expression: 1) retention of a single intron within *fgfr4* when maternal and zygotic TAF15 is depleted, and 2) reduction of total *fgfr4* transcript when zygotic TAF15 alone is depleted. The two mechanisms of gene regulation (post-transcriptional vs transcriptional) suggest TAF15-mediated gene regulation is target and cofactor-dependent, depending on the milieu of factors that are present at different times of development.

## INTRODUCTION

The FET family of atypical RNA-binding proteins includes Fused in sarcoma (Fus), Ewing’s sarcoma (EWS), and the TATA-binding protein-associate factor 15 (TAF15). This is a family of heterogeneous nuclear ribonuclear particle (hnRNP) proteins that contain domains for transcriptional activation, RNA binding, and DNA binding (Schwartz, Cech, & Parker, 2015). FET family proteins function in both RNA Polymerase II-mediated transcription and pre-mRNA splicing (Schwartz et al., 2015; Tan & Manley, 2009). Among vertebrates, the three FET members are highly conserved from fish to mammals, suggesting an independent and specialized requirement for each protein (Schwartz et al., 2015). FET proteins have been investigated primarily as components of fusion oncogenes; following abnormal chromosomal translocations, FET protein N-terminal low-complexity/activation domains are found fused to various DNA-binding proteins, contributing to the formation of various cancers (e.g. sarcomas and leukemias) as well as neuronal degenerative diseases (Crozat, Åman, Mandahl, & Ron, 1993; Delattre et al., 1992; King, Gitler, & Shorter, 2012; Kovar, 2011; Martini et al., 2002; Neumann et al., 2011; Panagopoulos et al., 1999; Rabbitts, Forster, Larson, & Nathan, 1993; Sjögren, Meis-Kindblom, Kindblom, Åman, & Stenman, 1999; Tan & Manley, 2009; Vance et al., 2009). It has only been more recently that the functions of these proteins have been examined in their full length, “wild-type”, form (Dichmann & Harland, 2012; Schwartz et al., 2015; Tan & Manley, 2009). Studies of the structural, functional, and biochemical properties of the FET family proteins determined that these proteins have multiple functions, such that FET proteins may have evolved to facilitate the complex coupling of transcription and mRNA processing that occurs in multicellular organisms (Kato et al., 2012; Schwartz et al., 2015; Schwartz, Wang, Podell, & Cech, 2013).

The majority of work that has contributed to our understanding of FET protein biology and disease mechanism has been carried out in cell lines and mouse models (Hicks et al., 2000; Li et al., 2007; Scekic-Zahirovic et al., 2016; Sharma et al., 2016; Svetoni et al., 2016; Kapeli et al., 2016) with little known of the role of FET proteins in embryonic development. Previous work from our lab examining the role of Fus in *Xenopus* development found that embryos depleted of Fus exhibit mesoderm differentiation defects and epithelial dissociation (Dichmann & Harland, 2012). The underlying mechanism of these phenotypes was retention of all introns in *fibroblast growth factor 8* (*fgf8*), *fibroblast growth factor receptor 2* (*fgfr2*), and *cadherin 1* (*cdh1*) transcripts (Dichmann & Harland, 2012). This study therefore showed that Fus is required for processing of a subset of transcripts in *Xenopus* development. Given the important role of FUS in *Xenopus* development, the perplexing potential for functional redundancy of FET family members in mouse (while remaining highly conserved throughout vertebrates), and the lack of basic research on FET protein functions, we examined the role of TAF15 in early *Xenopus* development; including the role of maternal versus zygotic TAF15.

To determine the role of TAF15 in early *Xenopus* development, we used RNA-sequencing (RNAseq) from single embryos depleted of maternal (M) and zygotic (Z) TAF15, using reagents that target all mRNA by inhibiting translation with Morpholino oligonucleotides, or just zygotic function using splice blocking MOs or CRISPR mediated mutagenesis. Upon evaluating the transcriptional changes that result from M+Z versus Z-only TAF15 depletion, we find a subset of target genes whose expression is regulated either post-transcriptionally (via intron retention) or by transcript level, depending on whether maternal or zygotic TAF15 is depleted. These results suggest that maternal TAF15 translation is limiting for splicing of a subset of mRNAS. Further, we show that during the time of zygotic genome activation, zygotic TAF15 modulates the expression of nascent target genes, acting at the transcriptional (rather than post-transcriptional) level. Interestingly, we find that in at least one case that we examined closely, maternal and zygotic TAF15 have a shared target gene (*fgfr4*), but that each act to regulate the target gene expression through post-transcriptional vs. transcriptional mechanisms, respectively. Here, we describe our findings as an example in *Xenopus* where the gene product TAF15: 1) uses distinct molecular mechanisms to regulate the expression of the same gene target (*fgfr4*) depending on the time of development in which TAF15 is expressed (maternal versus zygotic) and 2) ensures proper dorsoanterior neural development through two distinct molecular pathways (*fgfr4* and *ventx2*.*1*).

## METHODS

### Ethics statement

This study was carried out in strict accordance with the recommendations in the Guide for the Care and Use of Laboratory Animals of the National Institutes of Health. The protocol was approved by the Animal Care and Use Committee at the University of California, Berkeley.

### General Xenopus Embryo Culture

*Xenopus tropicalis* embryos were obtained through natural matings. For next day (daytime) matings, males were housed individually, and females were housed together, in four liter Rubbermaid® containers filled with two liters of water collected from the *X. tropicalis* housing racks. The night before the natural mating, males were boosted with 100 units (U) of human chorionic gonadotropin (HCG: Chorulon®, Merck, NADA NO.140-927, Code No. 133754) and females were primed with 10U HCG. The morning of mating, females were boosted with 200U HCG and paired with males. *X. tropicalis* embryos were collected using a disposable polyethylene transfer pipet (Fisherbrand®, Cat No. 13-711-7M), with the tip cut off to enlarge the opening. Embryos were dejellied and cultured as previously described (Khokha et al., 2002).

*Xenopus tropicalis* embryos were allowed to develop in 1/9X Marc’s Modified Ringer (MMR), until desired stage according to the normal table of development (Nieuwkoop & Faber, 1994).

### Whole-mount RNA in situ Hybridization

*Xenopus* embryos were fixed in MEMFA (0.1M MOPS pH7.4, 2mM EGTA, 1mM MgSO4, 3.7% v/v Formaldehyde) as previously described (Sive et al., 2000). RNA probes were labelled with digoxigenin-UTP, and chromogenic reactions were carried out by incubating hybridized embryos in Anti-Digoxigenin-AP Fab fragments, 1:3000 (Roche, 11 093 274 910), and the alkaline phosphatase substrate BM purpled (Roche, 11 442 074 001), as previously described (Sive et al., 2000). *Xenopus tropicalis* embryos were incubated in prehybridization buffer for at least one hour.

### Western Blotting

*Xenopus tropicalis* embryos were lysed in 20mM Tris–HCl pH 8.0, 50mM NaCl, 2mM EDTA, with 1x protease inhibitor (Roche cOmplete, Mini, EDTA-free, Product#11836170001), with 20uL per embryo 1% Triton™ X 100 detergent (freshly added ;Sigma T8787), homogenized by pipetting and freezing at -80°C. To pellet debris, lysates were spun at 2655 RCF (eppendorf Centrifuge 5417C) at 4°, and supernatant was transferred to new tube. To clear embryos of yolk, lysates were spun at 2655 RCF an additional two to three times, each time using a vacuum with a non-filtered p200 tip to briskly remove the yolk from top of the lysate. Lysate protein concentrations were measured with Bradford assays (Bio-Rad Protein Assay Dye Reagent Concentrate, Cat#500-0006) and read using a Molecular Devices, Spectramax M2 plate reader. Lysates were aliquoted for use and 6x loading dye was added. Samples were heated at 80°C for 10 minutes. Lysates were run on 8% polyacrylamide gel and run at 120V for 1 hour and forty-five minutes, eliminating proteins of 10-20 kDa. Proteins were transferred from gels using the semi-dry transfer system (BioRad Trans-Blot® SD Semi-Dry Electrophoretic Transfer Cell #170-3940) to Immobilon®-FL transfer membranes, PVDF (Millipore, IPFL00010), and blocked for one hour at room temperature with 1X Odyssey® Blocking Buffer (PBS) (LI-COR, 927-40000). Anti-TAF15 (TAFII68) antibody (Bethyl Laboratories, A300-309A) was used at 1:3000, anti-FGFR4 (CD334) antibody (Thermo Scientific, PA5-28175) was used at 1:2000, and anti-β-actin antibody (GeneTex, clone GT5512, GTX629630) was used at 1:5000, diluted in 5% BSA in TBS-tween (TBS-T), and incubated overnight at 4°C. Fluorescent secondary antibodies, Alexa Fluor® 680 goat anti-Rabbit IgG (Invitrogen, A-21109), IRDye® 800CW Donkey anti-Mouse IgG (LI-COR, 925-32212), were incubated at 1:10,000 for one hour at room temperature in the dark. Western blots were visualized and quantified using a LI-COR imager and software (LI-COR, Odyssey).

### Microinjection of Morpholino Antisense Oligonucleotides: Maternal and/or Zygotic TAF15 Depletion

Morpholino antisense and mismatch oligos (MOs) were designed and ordered from GeneTools LLC.

*taf15* translation-blocking (Maternal and Zygotic) MO:

5’-AGCTACTGGGATCTGAAGACATGAT-3’;

*taf15* splice-blocking (Zygotic only) MO: 5’-TTCCAAAACCTACCTTTGTTGCTGC-3’; Mismatch MO: ‘5-AGCTAGTCGCATCTCAACACATGAT-3’.

MOs were dissolved in nuclease-free water to 8.5ng/nL (1mM). Translation-blocking (17ng/cell), splice-blocking (8ng/cell), or mismatch (17ng/cell) MOs were injected into either one of two, or two of two cells, to deplete target mRNA from half or the whole embryo, respectively. To trace which cells contain MO, each MO was coinjected with the fluorescein-conjugated standard control oligo (GeneTools).

### RNA-extraction

RNA from single *Xenopus tropicalis* embryos was isolated using the Trizol® Reagent (Ambion Ref# 15596026), optimized for extracting small amounts of RNA from single *Xenopus tropicalis* embryos. Single embryos were collected in 200μl of Trizol® Reagent (Ambion Ref# 15596026), homogenized by pipetting, and stored at -80°C for a minimum of fourteen hours. Homogenized samples were thawed and incubate at room temperature (RT) for five minutes to allow complete dissociation of the nucleoprotein complex. 40uL of chloroform was added to each sample, vortexed, and incubated at RT for 2-3 minutes. Samples were centrifuged at 12,000 RCF for 15 minutes at 4°C and 90uL of the upper aqueous phase was removed and placed in new tube. RNA was precipitated with 100% isopropanol with 5ug of linear acrylamide (Ambion, AM9520) for pellet detection. RNA pellets were washed with 75% ethanol and air dried at RT for 5-10 minutes. RNA pellets were resuspended in 250uL of Milli-Q water and gently pipetted up and down and vortex. A second RNA extraction was performed by adding 250uL of Acid-Phenol:Chloroform, pH 4.5 (with IAA, 125:24:1) (Ambion, Cat#AM9720) following the manufacturer’s protocol. RNA was precipitated by adding 20uL 5M NH4OAC (Ammonium Acetate) (Ambion AM9070G) and 220uL of 100% isopropanol and RNA washed two times with 75% ethanol. RNA concentrations were measured using a nanodrop (Nanodrop ND-1000 Spectrophotometer) and used for RNAseq library preparation or qRT-PCR.

### RNA sequencing Library preparation and Analysis

Each paired-end library for RNA-sequencing (RNA-seq) was prepared using RNA extracted from single *Xenopus tropicalis* embryos; each condition is composed of three to four independently sequenced embryos. RNA-seq libraries were made strictly following the Low Sample (LS) Protocol from TruSeq RNA Sample Preparation v2 Guide. The important modification made to this protocol was that all reagents were used at half volume, except for the Bead Washing Buffer.

Final RNA-seq libraries were quantified using the KAPA Library Quantification Kits for Illumina sequencing platforms (Roche, Kit code KK4824). 100bp paired-end sequencing reads (Illumina HiSeq2000) were aligned to the *Xenopus tropicalis* genome version 7.1, and an annotation from Darwin Dichmann (not published). RNA-seq data analysis for differential gene expression was performed using both the Tuxedo Suite (Tophat, Bowtie, Cufflinks, Cuffdiff) (Trapnell et al., 2012) and the Bioconductor package DESeq (Love, Huber, & Anders, 2014). RNA-seq data analysis for differential intron-exon usage was performed using the Bioconductor package, DEXseq (Reyes, Anders, & Huber, 2014) (www.bioconductor.org). In brief, read counts were normalized to library size per feature. The p-values are the result of the statistical modeling peformed by DESeq2/DEXSeq and indicates to which degree the difference in expression of a given gene or exon is significant (in morphants compared to controls). The p-values were then adjusted for false discovery rates (also performed by DESeq2/DEXSeq) for the number of features since false positive calls are problematic when sampling a large number of features (genes or exons) as is done in gene expression studies.

RNA-seq alignments were visualized using the Integrative Genomics Viewer (IGV) from the Broad Institute (www.broadinstitute.org/igv/).

### Complementary DNA (cDNA) Synthesis and qRT-PCR

RNA was isolated from single *Xenopus tropicalis* embryos as described above. cDNA was synthesized using the iScript reverse transcription (RT) reaction protocol (BIO-RAD Cat#170-8841). Optimally, 1ug of total RNA was reverse transcribed, but total RNA yield from a single *Xenopus tropicalis* embryo varied. Therefore, all samples were normalized to ensure all RT samples contained the same volume and concentration of RNA. For each RT reaction, 7.5uL of 5x iScript RT Supermix was added to 30uL of RNA/H2O. For no RT (NRT) samples, RNA samples with yields below a usable limit were pooled and the sample volume was brought up to 30uL, and 7.5uL of 5x iScript Supermix no RT was added.

cDNA samples were diluted to a working concentration of 5ng/uL and qRT-PCR reactions were performed with the SsoAdvanced SYBR Green Supermix (BIO-RAD Cat#172-5261) using a CFX96 Thermal Cycler (BIO-RAD) following the manufacturer’s suggested protocol. NRT and no template control (NTC) samples were included for each gene target.

Primer set pairs used in these studies:

***eef1a1***: F 5’-CCCTGCTGGAAGCTCTTGAC-3’;

R 5’-GGACACCAGTCTCCACACGA-3’

***fgfr4 intron 1***: F 5’-AGGGCTAAGCAGTGCCTGTA-3’;

R 5’-AATGCAAGAGCAGCTCCAAT-3’

***fgfr4 total***: F 5’-AGCCAGGAATGTTCTTGTGG-3’;

R 5’-TCCCATGTCAACACTCCAAA-3’

***ventx2***.***1***: F 5’-AACAGCCAGCTGTCTCCAGT-3’;

R 5’-GCTGTGTCCCTGTGTAGCAA-3’

***engrailed 2***: F 5’-GAAGACGACGACGACTGTCA-3’;

R 5’-AACTTTGCCTCCTCTGCTCA-3’

***bmp7***.***1 intron 3***: F 5’-TCCCCTCCTCTATGGCTTTT-3’;

R 5’-AGTGGTGCCCAAGATTCAAC-3’

***bmp7***.***1 total***: F 5’-CGGGAAAGGTTTGAAAATGA-3’;

R 5’-ATATCGAACACCAGCCATCC-3’

***cpl1 5’UTR***: F 5’-TATGCCTCCCCTGTGGATAG-3’;

R 5’-TACTTTGCCTGCCCTATGCT-3’

***cpl1 total***: F 5’-AGAGATCTGCCGCACTTTGT-3’;

R 5’-TGAACCACGTGGACCATAGA-3’

***dgka intron 9***: F 5’-GAGGGTGATGTGACCATGTG-3’;

R 5’-TCCATTTTAAGCCCAACAGC-3’

***dgka intron 11***: F 5’-GCGACCAATGAATGCTACCT-3’;

R 5’-TAAACATGCTGCTGGGTCAG-3’

***dgka total***: F 5’-TGGATGAGAGGTGGATGTGA-3’;

R 5’-CATCACACAGTGGGGATGAG-3’

***pdgfa intron 1***: F 5’-CCTCAGAGGCACTTTCCAAG-3’;

R 5’-TTGTGCTACAGAACCGCAAC-3’

***pdgfa total***: F 5’-CCAGAGAAGCGTTCTGTTCC-3’;

R 5’-ACACACGGAGGCCAGATTAG-3’

***per2 intron 1***: F 5’-GGCCTGTAATTTGGAGCTTTC-3’;

R 5’-CAAACCGGATGTGGGATTAT-3’

***per2 total***: F 5’-TGGACGGGAATCAAGAAAAG-3’;

R 5’-GAACCCTCTCAGCAAACAGC-3’

***rab15 intron 2***: F 5’-GGATGCTTTTGGTGGTGTTT-3’;

R 5’-ATTGGATGGTTTTGCCTCTG-3’

***rab15 total***: F 5’-GGATGAAGCTTGCAGAGGAG-3’;

R 5’-CTCCAGCTCCCTTTTGTGAG-3’

***srsf4 intron 5***: F 5’-TCTGATCCCCATTCAGTTGC-3’;

R 5’-GTTTTGCCTGCATAGCCAGT-3’

***srsf4 total***: F 5’-GAGCAAGGATAGGGACCACA-3’;

R 5’-TCTCTTTGCTACGGCTGGAT-3’

***zdhhc5 intron 6***: F 5’-GAAGCGGGCTATCTGAGTTG-3’;

R 5’-CCTTGGGGTCTGAGATTTGA-3’

***zdhhc5 total***: F 5’-ACTGGCCAATTTTAGCATGG-3’;

R 5’-AACCTGCACTCCCCTTACCT-3’

***gpr110 intron 9***: F 5’-GCCATTTAGTGATGGGCTGT-3’;

R 5’-CCAAGGCACACATACATGCT-3’

***gpr110 total***: F 5’-GCTTGTCATATCCCGTTGCT-3’;

R 5’-CTAGCCCACACGTAGCCATT-3’

***isl1 intron 3***: F 5’-CCCTCCTACCCTTTTCCAAG-3’;

R 5’-GGCTGAAGTGCGGAATCTAA-3’

***isl1 total***: F 5’-ACTTTGCCCTGCAGAGTGAC-3’;

R 5’-TGCCTCTATAGGGCTGGCTA-3’

Prior to use for quantitating transcripts, primer pair efficiency was tested on six standard controls (serial dilutions of 1/5): 25ng = 2.50E+04, 5ng = 5.00E+03, 1ng = 1.00E+03, 0.2ng = 2.00E+02, 0.04ng = 4.00E+01, 0.008ng = 8.00E+00.

### Glutaraldehyde Vibratome Sections

Following RNA *in situ* hybridization, embryos were selected for sectioning. Sectioning was performed as previously described (Young et al., 2014). Embryos were equilibrated in a PBS solution containing 20% sucrose, 30% BSA, 4.9% gelatin, and fixed with 1.5% glutaraldehyde. Embryos were mounted into a Peel-A-Way® disposable embedding mold (Polysciences, 18985). Once cured, the blocks were removed from the embedding molds and cut into sectioning prisms. Prisms were affixed to Heroscape dice (Hasbro) with super glue and mounted on a Pelco 101 Vibratome Series 1000 Sectioning System (Ted Pella, Inc.), sections were made with razor blades (Personna). 50-75micron sections were mounted on glass slides for imaging.

### Microinjection of CRISPR/cas9: Zygotic *taf15* Depletion

*X*.*tropicalis taf15* sgRNA was designed using the Giraldez Lab CRISPRscan (crisprscan.org). Sequence of *X*.*tropicalis taf15* sgRNA (including PAM): G<u>CTATGGTGGTTATGGAGG</u>AGG. sgRNA was generated by PCR amplification using the Phusion Polymerase (NEB, M0530S) following the manufacturer’s protocol. Forward primer CTAGCtaatacgactcactataGG<u>CTATGGTGGTTATGGAGG</u>gttttagagctagaa and reverse primer AAAAGCACCGACTCGGTGCCACTTTTTCAAGTTGATAACGGACTAGCCTTATTTTAACTTGCTATTTCTAGCTCTAAAC were used for sgRNA amplification. RNA *in vitro* transcription of sgRNA was performed using the T7 MEGAshortscript kit (Ambion/Life Technologies, AM1354) and sgRNA cleanup was performed using the MEGAclear kit (Ambion/Life Technologies, AM1908).

sgRNA injection cocktails included 1.5ng Cas9 protein, 400ng sgRNA, and Dextran555 (Life Technologies/Molecular Probes, D34679) diluted in water. Injection volume was 2nL. Embryos were injected in the two-cell embryo stage, injections were made into either one of two, or two of two cells, to deplete target mRNA from half or the whole embryo, respectively.

## RESULTS

### *taf15/*TAF15 is both maternally deposited, and zygotically transcribed, with a localized expression pattern

To study of the role of TAF15 in *Xenopus* development, we first determined the time and place of gene expression; both at the transcript and protein level (Figure 1). From the RNA *in situ* hybridization (ISH) data (Figure 1A), we observe that *taf15* is enriched in the animal hemisphere through cleavage and gastrula stages (Figure 1A). During neurulation, *taf15* changes from diffuse dorsal expression (Figure 1A, stage 13), to more specific expression around the neural plate but still throughout the ectoderm (Figure 1A, stage 15). Enrichment in the tailbud stage (Figure 1A, stage 26), is in dorsoanterior tissues of the embryo, particularly the brain, branchial arches, anlage of the ear and eye, and pronephros. Lastly, in the early tadpole (Figure 1A, stage 32), *taf15* is enriched in the central nervous system (anterior and posterior), branchial arches, otic vesicle, and the posterior domain of the eye. These data are consistent with protein abundance in Western blots: TAF15 is deposited maternally and present throughout embryogenesis, increasing in expression following zygotic genome activation (ZGA) (Figure 1B).

**Figure 1.**
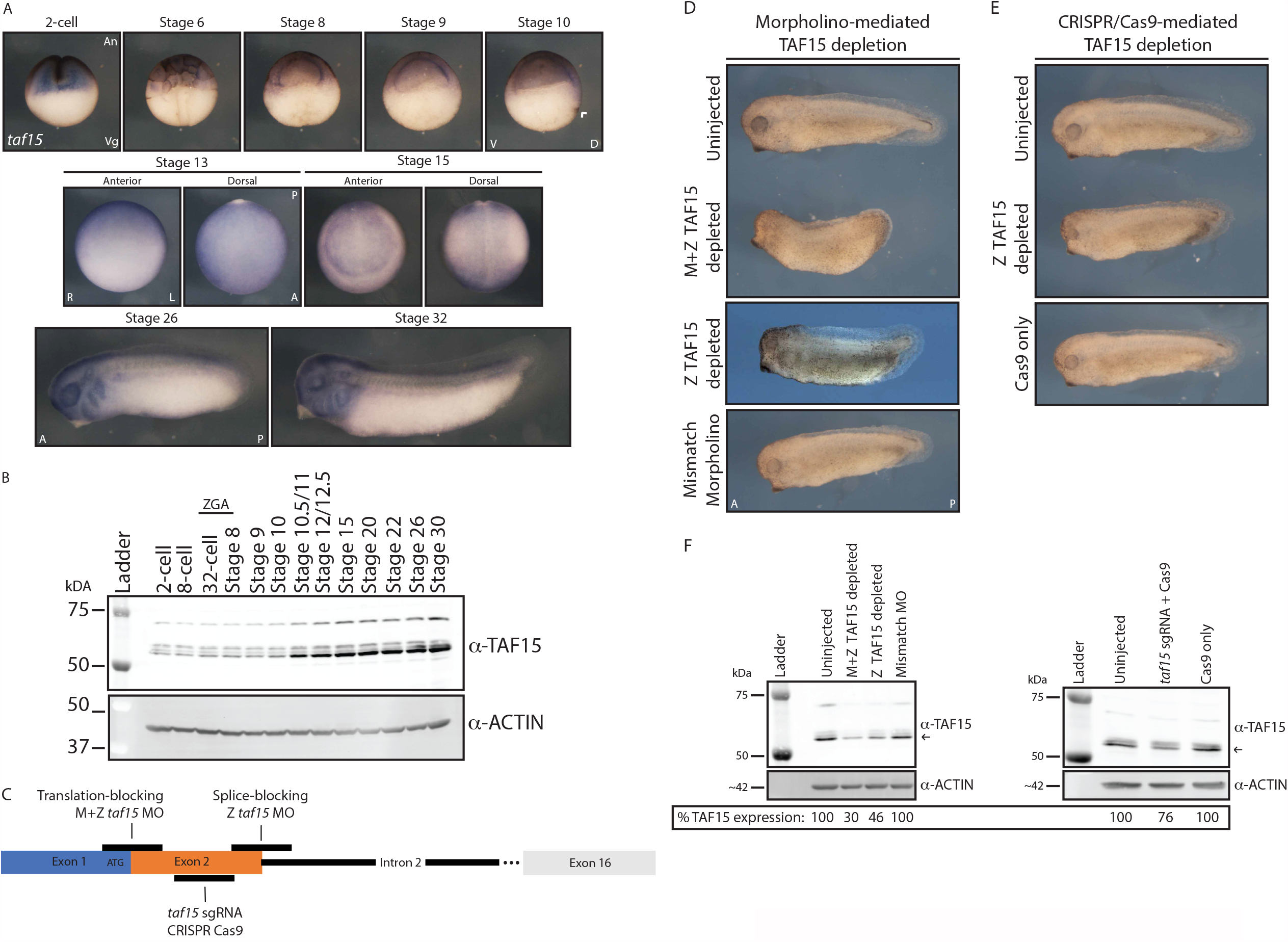
*taf15*/TAF15 expression and depletion phenotype. (A) Representative whole mount *in situ* RNA hybridization to *taf15* transcripts in *X. tropicalis* embryos. Stage 2-cell through stage 10; embryos were cut in half prior to hybridization. An = animal pole, Veg = vegetal pole, V = ventral, D = dorsal, Arrowhead = dorsal lip, R = right, L = left, A = anterior, P = posterior. (B) Western blot for TAF15 expression from two-cell through early tadpole; ZGA = zygotic genome activation. (C) Schematic indicating locations of morpholino and CRISPR/Cas9 target sites. (D) Translation-blocking (M+Z TAF15 depleted) and splice-blocking (Z TAF15 depleted) morpholino-mediated TAF15 depletion. A = anterior, P = posterior. (E) CRISPR/Cas9-mediated zygotic (Z)TAF15 depletion. A = anterior, P = posterior. (F) Representative Western blot for TAF15 protein expression in stage 15 embryos following morpholino or CRISPR/Cas9-mediated TAF15 depletion. Protein quantification is of the imaged blot using the most consistently expressed TAF15 band marked by an arrow, normalized to the corresponding ACTIN band; percent expression is relative to the uninjected condition; >3 blots were analyzed. (A) Representative images of >10 embryos. (D,E) Representative images of >12 embryos. (C,D,E) All injections for downstream imaging and Western blot analysis were into both cells of 2-cell stage embryos.

### *taf15*/TAF15 depletion leads to gross morphological defects

To elucidate the role of TAF15 in development, we depleted both maternal and zygotic (M+Z) TAF15 protein expression using a translation-blocking morpholino antisense oligonucleotide (MO) and zygotic (Z) expression alone using either a splice-blocking MO or CRISPR-associated protein-9 (Cas9) nuclease technology (CRISPR/Cas9) (Figure 1C-F). Following MO-mediated M+Z TAF15 depletion (by translation-blocking morpholino), embryos exhibit gross morphological defects including a shortened anterior-posterior axis, loss of dorsal and posterior fin structures, reduced eyes and dorso-anterior head structures (Figure 1D). Not all structures are defective, for example the cement gland appears unaffected (Figure 1D). These phenotypes are consistent with the *taf15* expression observed by ISH (Figure 1A, Stage 32). While morphological defects were clear in stage 32 tadpoles, early embryos appeared fairly normal, although gene expression changes were evident following TAF15 depletion (Figures 2-7). Embryos injected with a translation-blocking MO containing five mismatches to the *taf15* transcript (Mismatch) did not phenocopy the *taf15* morphants, suggesting the effects of the MO are specific to TAF15 depletion (Figure 1D). Following Z TAF15 depletion, either by splice-blocking MO or CRISPR/Cas9-mutagenesis, embryos exhibit a milder phenotype compared to the M+Z TAF15 depletion, with the most consistent phenotypes between conditions being reduced eyes and dorso-anterior head structures (Figure 1D-E). Importantly, embryos injected with Cas9 protein alone (Cas9 only) were similar to uninjected embryos, again suggesting that *taf15* sgRNA+Cas9 phenotype is specific to the guide that mediates TAF15 depletion (Figure 1E). The discrepancy in the phenotypes observed following M+Z or Z only TAF15 depletion suggests some separable developmental roles for maternal and zygotic TAF15, as explored further below.

**Figure 2.**
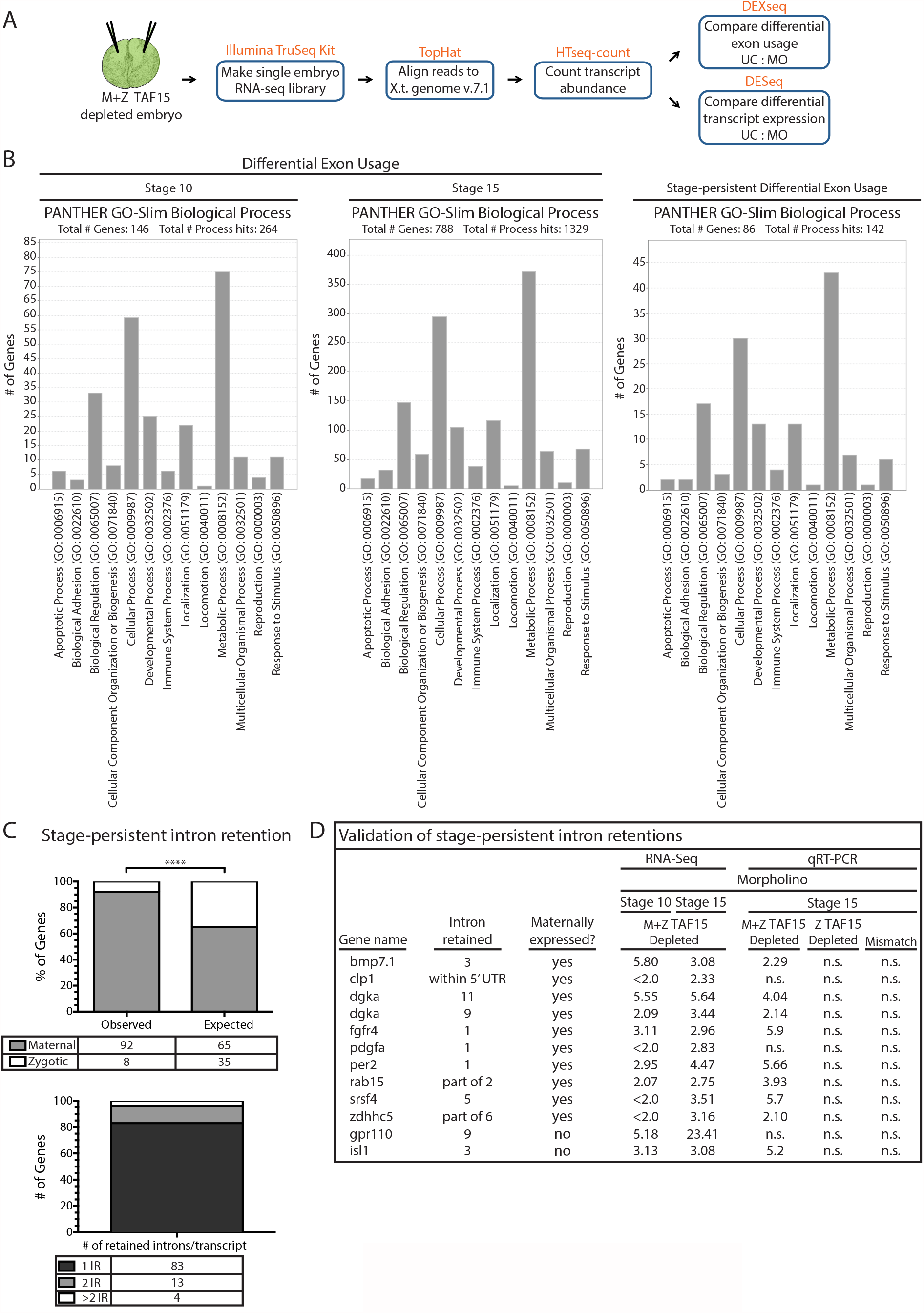
TAF15 depletion by translation-blocking morpholino leads to intron retention. (A) Schematic for maternal and zygotic TAF15 depletion via injection of translation-blocking morpholino and subsequent single embryo RNA-sequencing analysis. (B) PANTHER GO-Slim biological process classification of genes with differential exon usage following maternal and zygotic TAF15 depletion. (C) Genes with intron retention at both stages 10 and 15 classified by their maternal or zygotic expression, as well as the number of introns retained per transcript. **** P value = 1.50723E-08. IR = intron retention. (D) qRT-PCR validation of RNA-seq results for genes with intron retention at both stages 10 and 15; qRT-PCR results expressed in fold change relative to uninjected controls; intron expression normalized to *eef1a1* and to respective total gene expression; n = 3 individual embryos. n.s. = not significant. All injections for downstream RNA sequencing and qRT-PCR analysis were into both cells of 2-cell stage embryos.

To evaluate depletion efficiency, Western blot analysis was used to measure total TAF15 following MO or sgRNA+Cas9 injection (Figure 1F). One possible reason for a milder phenotype following Z TAF15 depletion (either MO or CRISPR/Cas9-mediated) is likely the reduced TAF15 depletion efficiency in these conditions, as compared to M+Z TAF15 depletion (Figure 1D,E,F). Indeed, none of the treatments eliminated TAF15 completely, illustrating the sensitivity of development to the dose of TAF15, confirmed below by RNA sequencing (RNAseq). While we attempted to rescue the phenotypes with injected *taf15* mRNA, overexpression is also teratogenic, and we were unable to find a dose that gave robust rescue. This is consistent with other experiments where both increased and decreased expression of splicing regulators induces developmental defects (Dichmann, Fletcher, & Harland, 2008; Iwasaki & Thomsen, 2014). Multiple protein bands (most prominently at ∼55kDa and ∼60-65kDa) are observed for TAF15 by Western blot (Figure 1A,E and Figure 4F). There are two known isoforms of *taf15* in *Xenopus tropicalis*. XM_012956707.2 comprises 1% of the *taf15* transcript and NM_001004806.1 comprises the remaining 99%; all of our depletion tools target the latter 99%. TAF15 depletion affects all protein bands, suggesting they are specific to TAF15 and targets of our depletion tools. As such, we suggest that multiple bands arise from post-translational modifications.

### Maternal TAF15 regulates splicing of developmental regulators

In addition to the gross morphological defects that follow TAF15 depletion, changes in gene expression were analyzed by single-embryo RNA sequencing (RNAseq). RNAseq libraries were generated from stage 10 (gastrula stage) and 15 (neurula stage) embryos injected with TAF15 translation-blocking MO (resulting in M+Z TAF15 depletion) (Figure 1C and 2A). The translation-blocking morphants were selected for sequencing as these embryos displayed a more severe depletion phenotype (Figure 1D,E) and indicated a consistently more robust TAF15 depletion (as assayed by Western blot Figure 1F). TAF15 is a member of the FET family of proteins and the family member Fused in liposarcoma (Fus) is necessary for the proper mRNA splicing of developmental regulators in *Xenopus* (Dichmann & Harland, 2012). To understand how widespread, or conserved, the roles of FET proteins are in splicing regulation in *Xenopus* we examined changes in intron/exon usage and gene expression levels using the bioconductor packages DEXseq and DESeq, respectively (Love, Huber, & Anders, 2014; Reyes, Anders, & Huber, 2014) following M+Z TAF15 depletion (Figure 2A).

Following DEXseq, a two-fold threshold cutoff was applied to the differential exon usage (DEU) gene candidates, followed by PANTHER GO-slim Biological Process analysis. In stage 10 embryos, 228 transcripts were found to exhibit DEU, and 1,429 in stage 15 embryos (Supplemental Tables 1 & 2); 146 and 788 of which were assigned a PANTHER GO-slim Biological Process respectively (Figure 2B). For the 100 genes with affected differential exon usage at both stages 10 and 15, termed “stage persistent” (Supplemental Table 3), we classified 86 genes with an identified PANTHER GO-slim Biological Process (Figure 2B). Importantly, because DEXseq measures exon usage, it is not an increase or decrease in gene expression that is measured in this “stage-persistent” DEXseq cohort, but instead genes with conserved differences in exon usage at both stages 10 and 15. Consistently we found enrichment for genes involved in cellular and metabolic processes across all stages (Figure 2B). Using a database designed to differentiate transcripts that are 1) “maternal”, meaning both maternally deposited and zygotically transcribed (present in the egg and expressed upon ZGA) (e.g. *fgfr4*) from 2) “zygotic”, transcripts that are exclusively zygotically transcribed (e.g. *isl1*), we were able to compare the percentage of observed target genes with DEUs that have maternal expression to the percentage of all *X. tropicalis* genes with maternal expression (database provided by the Rokhsar lab at U.C. Berkeley; not shown). The “observed” category is comprised of the 100 stage-persistent genes that show differential exon usage (Supplemental Table 3) and whether they have maternal and zygotic (“maternal”) or exclusively zygotic (“zygotic”) transcript expression. Here we observe 92% of these stage-persistent DEU target genes as transcripts known to have both maternal and zygotic expression; with the remaining 8% composed of exclusively zygotically expressed genes. The “expected” category represents the percentage of all annotated *X. tropicalis* that have maternal and zygotic (“maternal”) or exclusively zygotic (“zygotic”) transcript expression. Here we observe 65% of the *X. tropicalis* transcripts have both maternal and zygotic expression with the remaining 35% exclusively zygotically expressed (Figure 2C). We were therefore surprised to find 92% of the stage-persistent genes to be transcripts that are both maternally deposited and zygotically transcribed. These data suggest a preference for splice regulation of transcripts that are present throughout development (from egg through ZGA) in this M+Z TAF15 depletion condition, since only 65% of annotated *X. tropicalis* transcripts are both maternally deposited and zygotically transcribed (Figure 2C). Alternatively, it is also possible that intron retention within exclusively zygotic transcripts is less abundant than expected in the M+Z TAF15 depleted embryos. Interestingly, of the 100 stage-persistent genes, 83 exhibit one intron retention, 13 have two introns retained, and four have more than two introns retained (Figure 2C). Of the 100 “stage-persistent” DEU genes, we found 53 distributed throughout the top 100 DEU genes of the stage 10 embryo (as sorted by adjusted P value) suggesting that this approach of looking for conserved DEUs between stages 10 and 15 is robust for finding splicing events with an early and lasting effect throughout development. To ensure we further investigated genes with true DEUs, we used qRT-PCR to validate a subset of the DEU candidates that were found by RNAseq of TAF15-depleted embryos to have robustly retained introns (of which there were 29/100; Supplemental Table 3, column “Retained introns as visualized by DEXseq gene modeling”). 9/12 (75%) of these DEXseq RNA-seq results were validated by qRT-PCR (Figure 2D). These 12 cases of intron retention correspond to a total of 11 genes due to the validation of two introns belonging to *dgka*. Surprisingly, this qRT-PCR data lead to the finding that intron retention is exclusively found in embryos following M+Z TAF15 depletion; intron retention was never observed by qRT-PCR in embryos depleted of zygotic TAF15 or injected with mismatch MO (Figure 2D). Additionally, we show that the splice-blocking morpholino is specific to Z TAF15 depletion only and does not have an effect on maternally deposited TAF15 (Supplemental Figure 1); further supporting our hypothesis that splice changes are due to dose dependence of TAF15, which requires translation of maternal *taf15* mRNA.

### Depletion of M+Z TAF15 leads to intron retention in *fgfr4*

A two-fold expression cutoff was applied to the DEXseq and DEseq results (except for stage 10 DESeq) and the candidate genes were sorted by their adjusted P (padj) values. DEXseq results indicate *fgfr4* intron 1 to be the 1^st^ and 7^th^ hit in stage 10 and 15 embryos respectively (Supplemental tables 1 and 2). Numerous top-ranking candidate DEUs were visualized using the Integrative Genomics Viewer (IGV) and DEXseq DEU models at both stage 10 and stage 15 to gauge the significance of each DEU. Of these top candidates, *fgfr4* displayed the most robust intron retention (Supplemental figures 2 and 3). Additionally, FGFR4 was an intriguing candidate because it is known that FGFRs are alternatively spliced. Examining the levels of *fgfr4* expression, DESeq results indicate the *fgfr4* transcript to be the 20th and 133^rd^ hit in stage 15 and 10 embryos respectively (Supplemental tables 4 and 5). These results were validated by qRT-PCR in M+Z TAF15-depleted single embryos using *fgfr4* intron 1-specific primers; *fgfr4* intron 1 was assayed across all depletion conditions yet was not detected outside of the M+Z TAF15 depletion condition (Figure 3B). To confirm that *fgfr4* intron 1 qRT-PCR results were specific to retention (as suggested by visualization by IGV; Figure 3A) and not simply increased due to an overall upregulation of *fgfr4* pre-mRNA, the *fgfr4* intron 1 expression levels (*fgfr4* intron 1-specific primers, red primer set Figure 3A) were normalized to the total *fgfr4* expression levels (Figure 3B; *fgfr4* total transcript-specific primers, blue primer set Figure 3A). As a result of normalizing *fgfr4* intron 1 expression to total *fgfr4* expression, conditions that do not retain *fgfr4* intron 1 will have a relative expression equal to 1 within this analysis (Figure 3B). Consistent with our qRT-PCR findings in Figure 2D, intron retention is not observed following zygotic TAF15 depletion by morpholino or sgRNA/Cas9 (Figure 3B). Interestingly, however, loss of zygotic TAF15 (both by MO and CRISPR/Cas9) does lead to an overall reduction in total *fgfr4* expression (Figure 3C); these results contrast sharply with increased expression and intron retention of *fgfr4* expression following M+Z *taf15* depletion (Figure 3C). Importantly, *fgfr4* expression is unaffected following injection of control morpholino or Cas9 protein alone, suggesting that the changes in *fgfr4* expression are specific to TAF15 depletion (Figure 3B-C).

**Figure 3.**
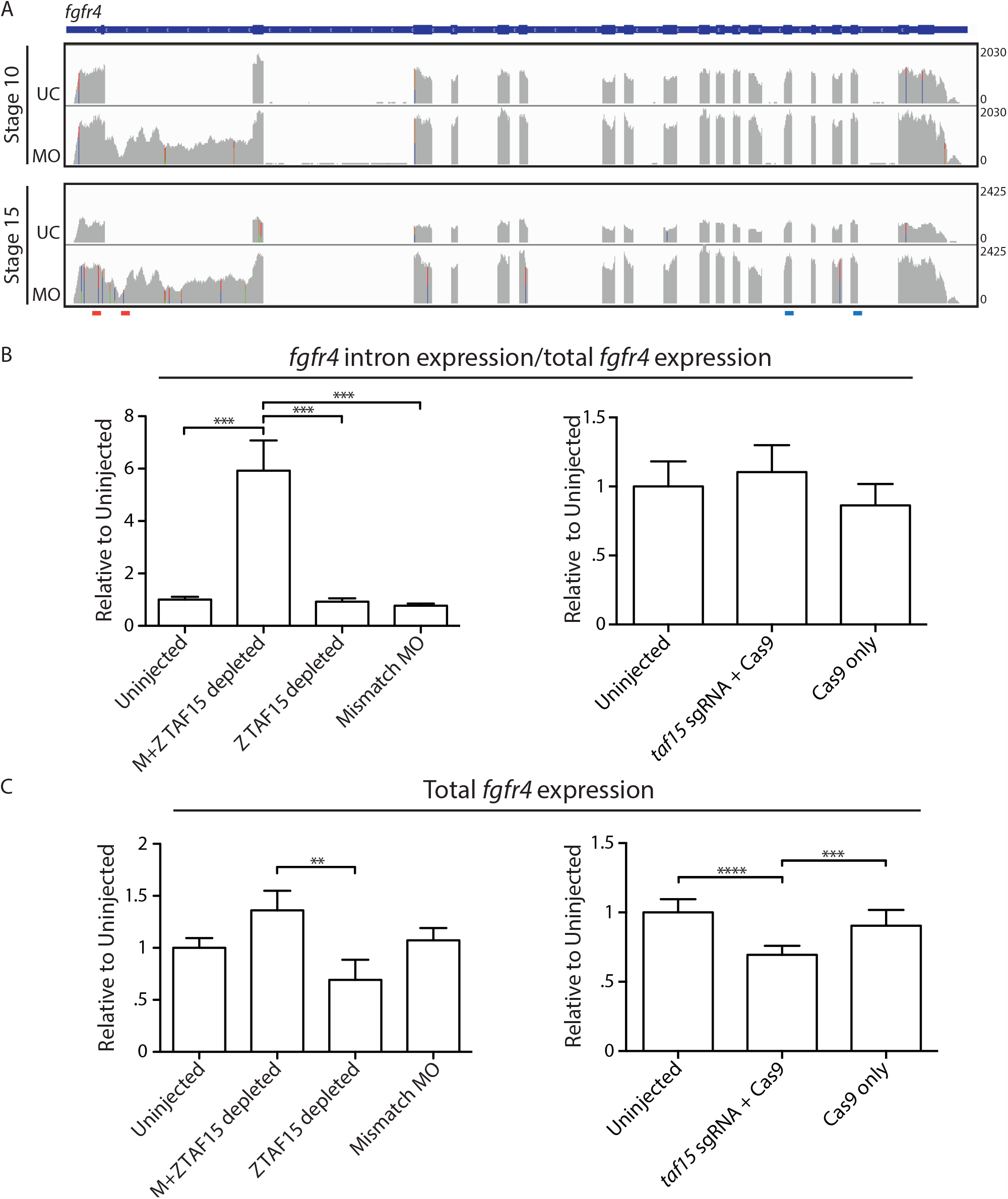
TAF15 depletion by translation-blocking morpholino leads to single intron retention in *fgfr4*. (A) Visualization of *fgfr4* RNA-seq reads with Integrative Genome Viewer aligned with gene model in blue. Red bars indicate qRT-PCR primers to measure retained intron. Blue bars indicated qRT-PCR primers to measure total transcript. UC = uninjected control; MO = M+Z TAF15-depleting morpholino (B) qRT-PCR for *fgfr4* intron 1 expression in stage 15 embryos. Intron expression levels are normalized to total *fgfr4* expression. *** P value = <0.0001. (C) qRT-PCR for total *fgfr4* expression on stage 15 embryos. ** P value = <0.005; *** P value = <0.001; **** P value = <0.0001. (B,C) Embryos treated as indicated on the *x*-axis; *y*-axis shows *fgfr4* expression relative to *eef1a1* and normalized to uninjected embryos. Error bars = standard deviation; n = 9 individual embryos. All means were compared by one-way ANOVA followed by Tukey post-hoc analyses. All injections for downstream qRT-PCR analysis were into both cells of 2-cell stage embryos.

**Figure 4.**
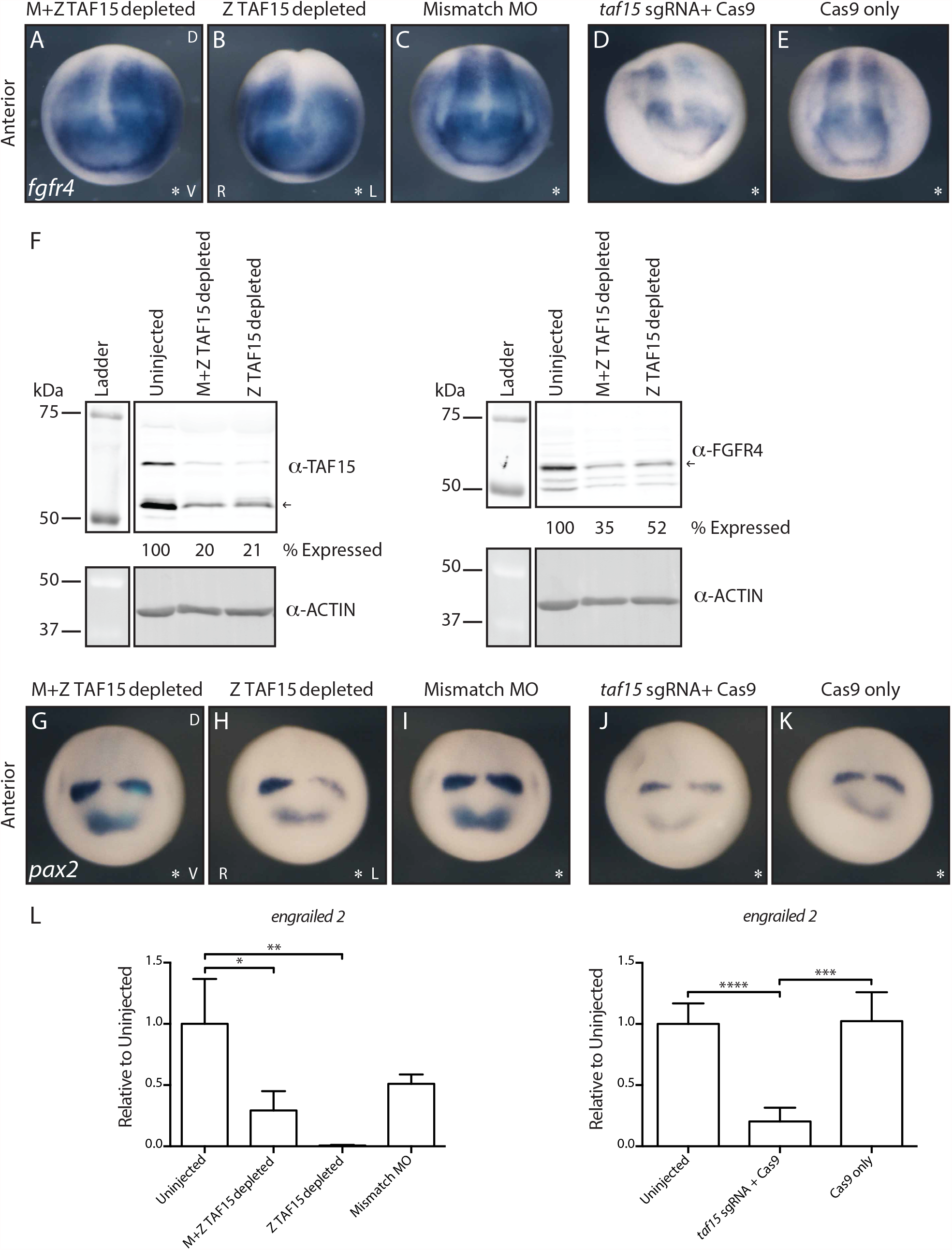
TAF15 depletion leads to downregulation of FGFR4 and downstream targets of FGFR4. (A-E) Representative whole mount *in situ* hybridization to *fgfr4* transcripts in TAF15-depleted *X. tropicalis* embryos; * = injected side; D = dorsal; V = ventral; R = right; L = left. (F) Representative Western blot for TAF15 and FGFR4 expression in stage 15 TAF15-depleted embryos. Protein quantification is of the imaged blot using the most consistently expressed TAF15 or FGFR4 band marked by an arrow, normalized to the corresponding ACTIN band; percent expression is relative to the uninjected condition; >3 blots were analyzed. (G-K) Representative whole mount *in situ* hybridization to *pax2* transcripts in TAF15-depleted *X. tropicalis* embryos; n=9/condition; D = dorsal; V = ventral; R = right; L = left. (L) qRT-PCR for *engrailed2* expression in stage 15 embryos treated as indicated on the *x*-axis; *y*-axis shows expression relative to *eef1a1* and normalized to uninjected embryos; n = 6 individual embryos. * P value = <0.05; ** P value = 0.002; *** P value = <0.001; **** P value = <0.0001; Error bars = standard deviation. All means were compared by one-way ANOVA followed by Tukey post-hoc analyses. (A-E, G-K) Representative images of >12 embryos. All injections for downstream RNA ISH were into one cell of 2-cell stage embryos; uninjected side serves as internal control. (F,L) All injections for downstream qRT-PCR analysis were into both cells of 2-cell stage embryos.

We next used RNA *in situ* hybridization (ISH) to total *fgfr4* to determine how the changes observed by qRT-PCR and RNAseq following TAF15 depletion may affect the expression pattern of *fgfr4*. M+Z depletion of TAF15 leads to an increase and expansion of the lateral domain of *fgfr4* expression just outside the neural plate (Figure 4A). We hypothesize that the intron-containing *fgfr4* transcripts fail to undergo degradation (necessitating normalization of intron levels to total transcript to avoid artificially inflating the amount of retained intron measured by qRT-PCR in Figure 3B) and are largely responsible for this diffuse increase in expression and that these transcripts are non-functional forms of *fgfr4*; indeed, M+Z TAF15 depletion results in reduced FGFR4 protein levels as measured by western blot (Figure 4F). Following zygotic depletion of TAF15 by MO or sgRNA/Cas9 we observe a decrease in *fgfr4* transcript and protein (assayed for morphants only) expression (Figure 4B, D, F). Importantly, *fgfr4* expression is unaffected following injection of control morpholino or Cas9 protein alone, suggesting that these results are specific to TAF15 depletion (Figure 4C, E). These results are consistent with the qRT-PCR data and suggest that the *fgfr4* transcripts are universally affected in each TAF15-depleted embryo, as the pattern of expression is largely unaffected (lateral expansion of *fgfr4* adjacent to the neural plate is observed in Z TAF15 depleted morphant embryos suggesting a modest delay in convergence as we observe complete neural tube closure). These findings further show that both M+Z and Z TAF15 MO depletion lead to a loss of FGFR4 protein (Figure 4F) and we hypothesize that this reduction is achieved through two different mechanisms: M+Z TAF15 depletion leads to intron retention in *fgfr4*, resulting in an early stop codon (data not shown), and in reduced FGFR4, whereas Z TAF15 depletion leads to the reduction of total *fgfr4* mRNA expression, also resulting in reduced FGFR4.

Lastly, after determining that total FGFR4 expression is reduced following both M+Z and Z-only TAF15 MO depletion, we next determined if downstream targets of FGFR4 were affected. Indeed, the two midbrain/hindbrain markers, *pax2* and *engrailed 2*, show reduction both by ISH or qRT-PCR across all TAF15 depletion conditions (Figure 4G-L) (Hongo, Kengaku, & Okamoto, 1999). These results indicate that M+Z and Z-only TAF15 depletion lead to the downregulation of FGFR4, but through independent mechanisms of transcriptional regulation.

We previously showed that 83% of “stage-persistent” DEUs are characterized by the retention of a single intron (Figure 2C); this is consistent with what is observed in *fgfr4*. A broader analysis determined if first introns are preferred targets or if *fgfr4* is an exception. In examining the top 10 genes with affected DEUs (as sorted by adjusted P value) at both stages 10 and 15, we found the single retained introns are not restricted to first introns, but are dispersed in different transcripts (Supplemental Figures 2 and 3; Supplemental Table 7). Therefore, retention of a single intron is the most common characteristic of DEU following M+Z TAF15 depletion (13/20 cases) within this cohort, contrasting with the effects of Fus depletion, which affects all introns of DEU genes.

### Measuring changes in gene expression following TAF15 depletion

It is unknown if and how the FET family of proteins regulates levels of gene expression in *Xenopus*. Our data thus far suggests that TAF15 controls both mRNA splicing and overall levels of a subset of RNAs in the developing embryo. Concurrent with analyzing how TAF15 depletion affects changes in intron/exon usage (DEXseq), we also examined levels of transcript abundance using DESeq (Figure 5).

**Figure 5.**
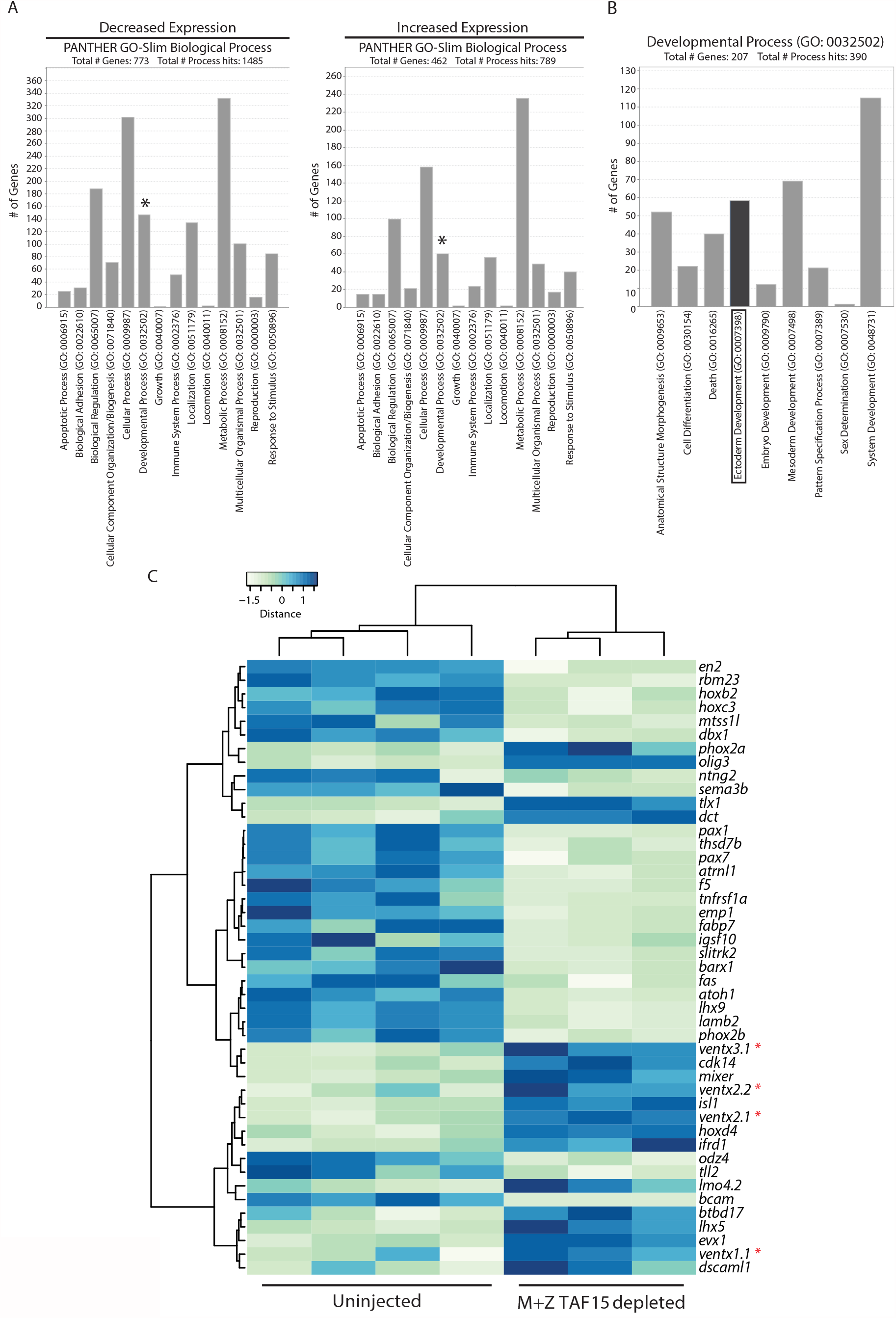
TAF15 depletion leads to upregulation of the *ventx* family of transcription factors. (A) PANTHER GO-Slim Biological Process classification of differentially expressed genes following maternal and zygotic TAF15 depletion; stage 15 *X. tropicalis* embryos. Asterisks mark Developmental Process genes combined into panel B. (B) Developmental Process subcategories. Boxed Ectoderm Development visualized in panel C. (C) Expression level heatmap of TAF15 target genes involved in Ectoderm development. Asterisks mark *ventx* family members. Columns represent RNAseq data from a single embryo.

Following DESeq, in an effort to focus on a few candidate genes that exhibit differential expression following TAF15 depletion, a two-fold threshold cutoff was applied to the differentially expressed gene candidates, followed by PANTHER GO-slim Biological Process analysis. In stage 15 embryos, 2,094 transcripts were found to exhibit a two-fold increase or decrease in gene expression (Supplemental Table 6), 1,235 of which were assigned a PANTHER GO-slim Biological Process (Figure 5A; 773 total genes with decreased expression plus 462 total genes with increased expression). Our *taf15* ISH data suggest that *taf15* has a specific expression pattern, therefore, we looked more closely at those genes that fall under the PANTHER GO-slim Developmental process (Figure 5A, asterisk); 207 genes were found to be differentially expressed within this classification (Figure 5B; combined Developmental Process results from Figure 5A). More specifically, our *taf15* ISH data suggests that *taf15* is expressed in the ectoderm at stage 15 (Figure 1A, Stage 15), therefore the subset of transcripts annotated with the Ectoderm development classification were more closely examined (Figure 5B, C). One family of genes in this category, that increases in expression following M+Z TAF15 depletion, is the VENT family of homeodomain transcription factors (*ventx*), suggesting a role for TAF15 suppression of *ventx* expression in the ectoderm (Figure 5C). Interestingly, *ventx* genes act in a positive feedback loop with the bone morphogenetic proteins (BMP), which specify the ventral domain of the Xenopus embryo (Onichtchouk, Glinka, & Niehrs, 1998; Sander et al., 2007). Because *taf15* is expressed in the future dorsal domain of the gastrula (Figure 1A), we were intrigued to find a gene family (*vent*) that functions in ventral tissue development, and is upregulated upon TAF15 depletion. This suggests an early and lasting regulatory relationship between *taf15* and *ventx*. Of the four *ventx* paralogs upregulated following M+Z TAF15 depletion, *ventx2*.*1* acts upstream of *ventx1*.*1, ventx2*.*2, and ventx3*.*1* therefore we investigate the relationship between *ventx2*.*1* and *taf15* further (Schuler-Metz, Knöchel, Kaufmann, & Knöchel, 2000).

### TAF15 regulates dorsoanterior development by repressing *ventx2*.*1*

To better understand the relationship between *taf15* and the *ventx* genes, we analyzed their expression patterns further. *In situ* hybridization revealed a subset of complementary expression patterns between *taf15* and *ventx2*.*1*, consistent with inhibition of *ventx2*.1 by *taf15* (Figure 6A-H). At stage 10, *taf15* is expressed in the embryonic domain that will give rise to future dorsal structures whereas *ventx2*.*1* expression is markedly absent from this region and is instead expressed in the embryonic domain that will give rise to future ventral structures (Figure 6A-D). *ventx2*.*1* and *taf15* continue their complementary expression into the neurula stage (Figure 6E-H). To visualize the extent of complementary expression, medial cross sections along the dorsoventral axis were made through stage 15 embryos following ISH. These sections reveal that *taf15* is strongly expressed throughout the neuroectoderm and in both the epithelial and sensorial layers of the ectoderm whereas *venx2*.*1* is strongly expressed in the underlying lateral plate mesoderm and only faintly expressed in a portion of the neuroectoderm (Figure 6E’, F’). One domain where *taf15* and *ventx2*.*1* exhibit overlapping expression is in the forebrain (Figure 6E-H).

**Figure 6.**
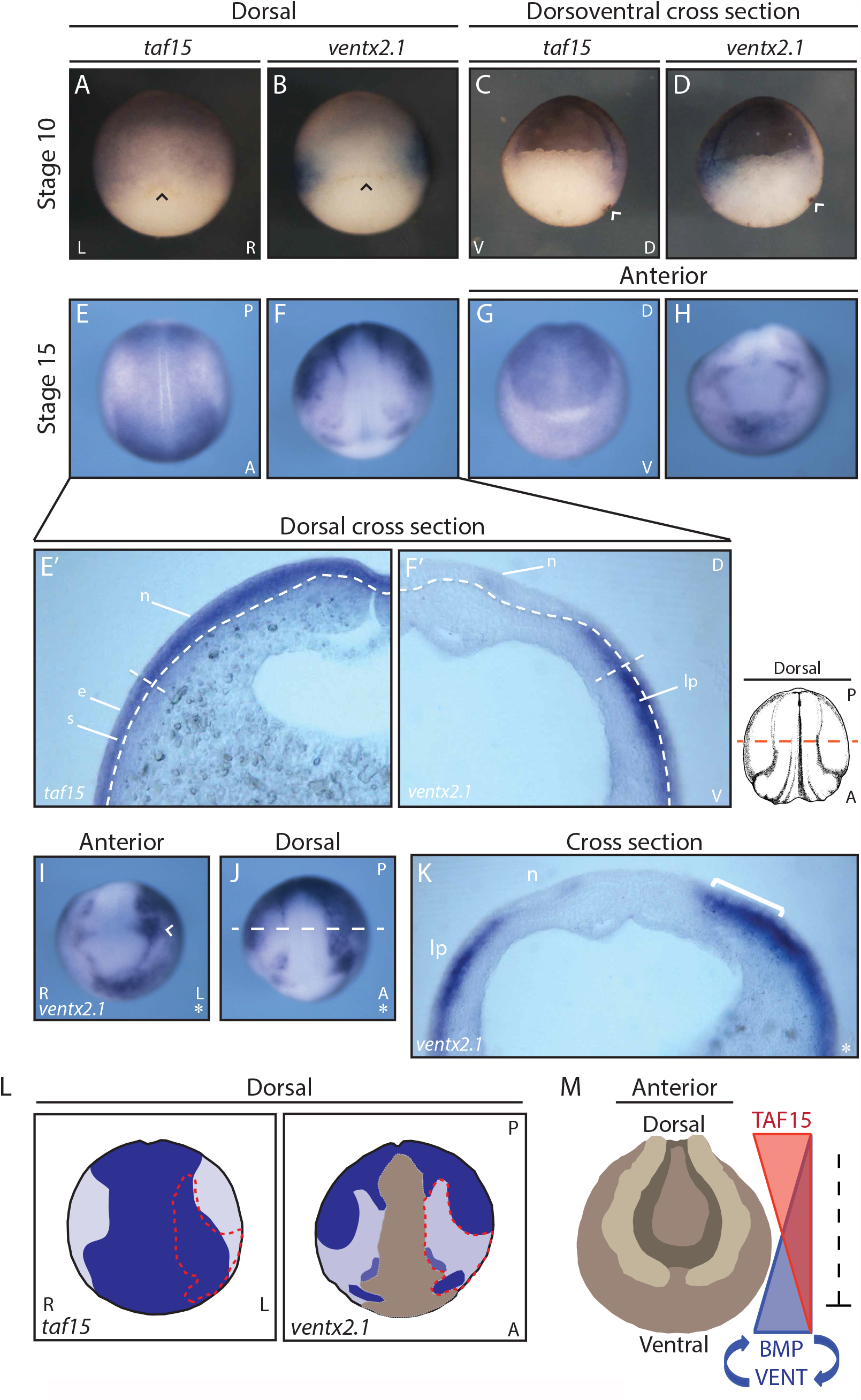
*taf15* and *ventx2*.1 exhibit complementary expression patterns and TAF15 depletion leads to expansion of *ventx2*.*1* expression into the neurectoderm. (A,C,E,G,E’) Whole mount *in situ* hybridization to *taf15*. (B,D,F,H,F’,I,J,K) Whole mount *in situ* hybridization to *ventx2*.*1*. (A-D) Arrowhead = dorsal lip. (E’,F’) White dotted lines = boundaries between the neural ectoderm and lateral plate mesoderm; red dotted line = site of cross-section. (I,J) Arrowhead = eye enlage; white dotted line = site of cross-section. (K) white bracket = expanded *ventx2*.*1* expression. (L) Schematic comparing *taf15* and *ventx2*.*1* expression. Dark blue = stronger expression; light purple = weaker expression; red dotted line = region of *ventx2*.*1* expansion. (M) Diagram of model for TAF15 repression of the ventrolateral BMP/Vent circuit. L = left; R = right; V = ventral; D = dorsal; A = anterior; P = posterior; s = sensorial layer of the epidermal ectoderm; e = epithelial layer of the epidermal ectoderm; n = neural ectoderm; lp = lateral plate mesoderm; * = injected side. (A-H) Representative images of >10 embryos. (E’,F’,K) Representative images of >5 cross-sectioned embryos. (I,J) Representative images of >12 embryos. (I,J,K) All injections for downstream RNA ISH were into one cell of 2-cell stage embryos; uninjected side serves as internal control.

Having established by RNAseq that *ventx2*.*1* expression increases with M+Z TAF15 depletion, and that *taf15* and *ventx2*.*1* have some complementary expression patterns, we examined *ventx2*.*1* expression *in situ* in TAF15-depleted embryos. Embryos with depleted M+Z TAF15 showed increased *ventx2*.*1* expression in regions that normally have strong *taf15* and weak *ventx2*.*1* expression, specifically below the lateral neuroectoderm and prospective epidermal region (Figure 6 E,F,J,K (white bracket) and L). Additionally, *ventx2*.*1* expression increases in the region of the forebrain where *taf15* and *ventx2*.*1* are coexpressed (Figure 6 G,H,I (arrowhead) and L). Taken together, these data suggest that TAF15 suppresses *ventx2*.*1*; and presumably does so indirectly, since the genes are expressed in adjacent germ layers. The increased level of *ventx2*.*1* is consistent with the gross phenotype of reduced head structures observed in older TAF15-depleted embryos (Figure 1D,E & Figure 6L, red outline).

### TAF15 regulates *ventx2*.*1* throughout early embryogenesis

RNAseq data show that M+Z TAF15 depletion results in increased *ventx2*.*1* expression by neurula stage 15 but not at the earlier gastrula stage 10 (<2 fold change) (Figure 7A). However, qRT-PCR results, show *ventx2*.*1* expression significantly increased at both stage 10 and 15 following M+Z TAF15 depletion (Figure 7B); consistent with the complementary *taf15* and *ventx2*.*1* expression patterns observed by ISH at stage 10 (Figure 6A-D). Following Z-only TAF15 depletion, we observe that *ventx2*.*1* expression is unaffected at stage 10 by qRT-PCR; however, by stage 15 the Z-only TAF15 depleted embryos phenocopy those that are M+Z TAF15 depleted (Figure 7B). We hypothesize that the effects of Z-only TAF15 depletion on embryonic development are not observed as early as stage 10 because zygotic genome transcription has only just begun (possibly not giving the zygotic genome-targeting MO enough time to act), furthermore, maternal TAF15 could still be present and able to suppress the transcription of *ventx2*.*1*. Embryos injected with a mismatch MO to the translation start site of *taf15* do not exhibit changes in *ventx2*.*1* expression at either stage, suggesting that the changes in gene expression observed in M+Z and Z-only TAF15 depleted embryos are specific to morpholino-mediated TAF15 depletion (Figure 7B, E, J).

**Figure 7.**
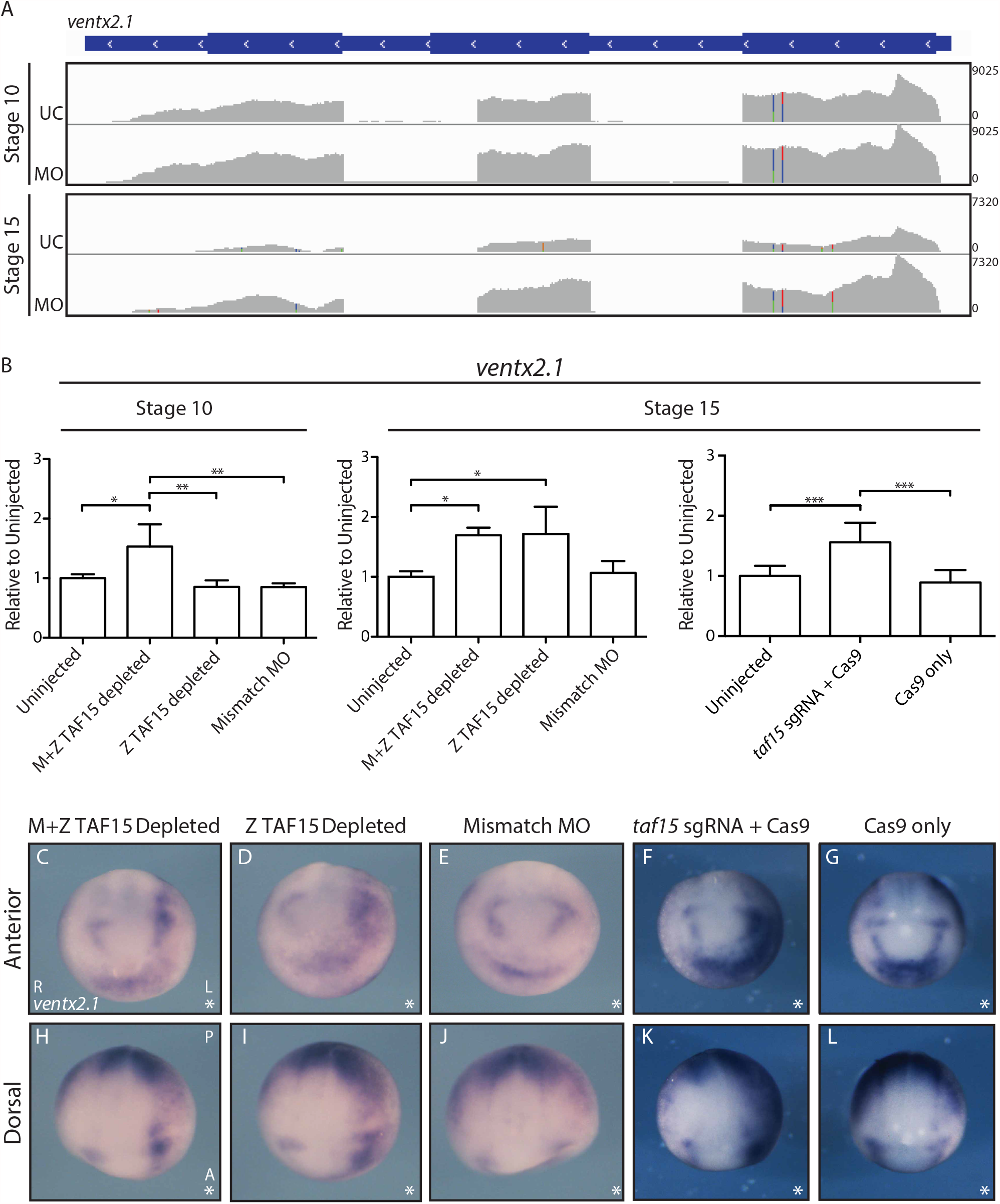
TAF15 depletion leads to increased and expanded *ventx2*.*1* expression. (A) Visualization of *ventx2*.*1* RNA-seq reads with Integrative Genome Viewer aligned with gene model in blue; UC = uninjected control; MO = M+Z TAF15-depleting morpholino. (B) qRT-PCR for *ventx2*.*1* expression in stage 10 and 15 embryos treated as indicated on the *x*-axis; *y*-axis shows expression relative to *eef1a1* and normalized to uninjected embryos; n = 9 individual embryos; * P value = <0.05; ** P value = <0.005; *** P value = <0.001; Error bars = standard deviation. All means were compared by one-way ANOVA followed by Tukey post-hoc analyses. All injections were into both cells of 2-cell stage embryos. (C-L) Representative whole mount *in situ* hybridization to *ventx2*.*1*; L = left; R = right; A = anterior; P = posterior; * = injected side. Representative images of >12 embryos. All injections for downstream RNA ISH were into one cell of 2-cell stage embryos; uninjected side serves as internal control.

To ensure that the increase in *ventx2*.*1* expression is not a morpholino-specific effect, we measured *ventx2*.*1* expression following TAF15 depletion with the CRISPR/Cas9 system. qRT-PCR data for *ventx2*.*1* expression in stage 15 sgRNA + Cas9-injected embryos phenocopies that observed in M+Z and Z-only TAF15 depleted embryos; importantly, injection of Cas9 alone does not affect *ventx2*.*1* expression level (Figure 7B). In addition to quantifying the changes in *ventx2*.*1* expression by qRT-PCR, the embryonic expression pattern of *ventx2*.*1* was also compared between MO and CRISPR/Cas9 conditions. We find that the expression pattern of *ventx2*.*1* following sgRNA *taf15 + Cas9* phenocopies that of M+Z and Z-only TAF15 depleted embryos (Figure 7C-L). Taken together, these qRT-PCR and ISH data suggest that changes in *ventx2*.*1* expression is specific to TAF15 depletion and that both maternal and zygotic TAF15 play a role in suppressing the expression of *ventx2*.*1* throughout development.

## DISCUSSION

Previous to our investigation, few studies have focused on the role of TATA-binding protein associated factor 15 (TAF15) in development. TAF15 is not considered a canonical TATA-binding protein associated factor (TAF) as it is not associated with all human TFIID complexes and has no ortholog in invertebrate species (Ballarino et al., 2012). However, it is this non-ubiquitous association with the core transcriptional machinery that interests us as this supports the hypothesis that TAF15 may be selective in the transcripts it regulates and, as a result, may have specific roles in development. TAF15 would not be the first TAF to be shown to have a specific role in development, for example, TAF3 is required for endoderm lineage differentiation and preventing the premature specification of neurectoderm and mesoderm in embryonic stem cells (Liu et al., 2011).

The analysis presented here demonstrates that TAF15 is deposited maternally in *Xenopus* eggs and is later expressed zygotically. We further show that TAF15 has an enriched expression pattern within the developing embryo with maternal *taf15* preferentially in the animal hemisphere and zygotic *taf15* enriched dorsally during gastrulation, throughout the neural ectoderm during neurulation, and in dorsoanterior tissues during tailbud and tadpole stages. Consistent with the expression pattern of *taf15* we find that embryos depleted of *taf15* have defects in head structures such as reduced fore/midbrain and eyes by the early tadpole stage. Interestingly, by employing RNA sequencing (RNAseq) to understand the role of TAF15 in development we made the surprising discovery that maternal and zygotic TAF15 exhibit different mechanisms by which they regulate FGFR4 expression: the depletion of maternal and zygotic TAF15 together downregulates FGFR4 expression at the post-transcriptional level via the retention of a single *fgfr4* intron, while depletion of zygotic TAF15 alone downregulates FGFR4 expression through the reduction of total *fgfr4* transcript. Additionally, we find that TAF15 is required to repress *ventx2*.*1* from dorsal and neural ectodermal tissues and that *taf15* and *ventx2*.*1* exhibit a complementary expression pattern through gastrulation and neurulation. The data presented here suggest TAF15 plays an integral and pleiotropic role in development of the dorsoanterior neural tissues and further suggest that the mechanism of gene regulation by TAF15 is target-dependent and subject to the milieu of factors that are present at different times of development, likely due to the presence of specific co-factors required for activity (Figure 8).

**Figure 8.**
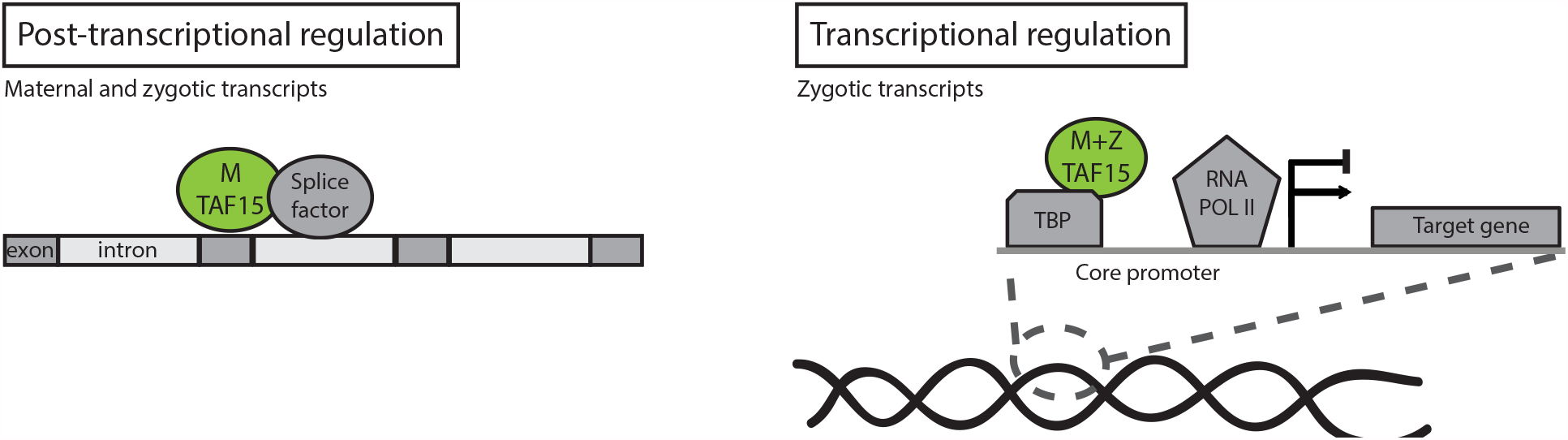
Model of TAF15 pre- and post-transcriptional regulation. M TAF15 = maternal TAF15; M+Z TAF15 = maternal and zygotic TAF15; TBP = TATA-binding protein; RNA POL II = RNA polymerase 2.

It is not unexpected that TAF15 plays a pleiotropic role in *Xenopus* development as FET proteins are associated with regulating numerous cellular activities including: cell proliferation, cell cycling, cell death, transcription, splicing, microRNA processing, RNA-transport, signaling, and maintenance of genomic integrity (Andersson et al., 2008; Ballarino et al., 2012; Gregory et al., 2004; Shiohama, Sasaki, Noda, Minoshima, & Shimizu, 2007). Furthermore, we expected that splicing could be a shared mechanism of gene regulation between Fus and TAF15 in *Xenopus* development as studies using photoactivatable ribonucleoside-enhanced cross-linking and immunoprecipitation (PAR-CLIP) found that FET proteins predominantly bind to intronic regions as well as the 3’UTR of genes (Hoell et al., 2011). However, we are surprised by how differently the two FET family members, Fus and TAF15, affect *Xenopus* development; embryos depleted of Fus fail to undergo gastrulation, due to the retention of all introns in a subset of target genes required for mesoderm differentiation and epithelial adhesion (Dichmann & Harland, 2012), whereas embryos depleted of TAF15 survive into the tadpole stage and exhibit defects specific to dorsoanterior head development due in part to dysregulation of *fgfr4* and *ventx2*.*1*, and in contrast to Fus, TAF15 depletion affects a subset of specific introns.

TAF15 provides an example of a gene product with two different mechanisms (post-transcriptional or transcriptional) by which to regulate the expression of the same target gene (FGFR4). Furthermore, we have shown that the mode of regulation is dependent on whether TAF15 is maternally deposited or zygotically transcribed. While there are classic examples of genes having separable maternal and zygotic developmental roles (e.g. β-catenin) (Heasman et al., 1994; Heasman et al., 2000), we are unaware of a case where the mechanism of regulatory action changes while the target remains the same.

Classical RNA splicing can be separated in two functional groups, constitutive and alternative splicing. Constitutive splicing is the process by which introns are removed (spliced), stitching together exons in the same order that they are found in the genome, producing one gene product (Boutz, Bhutkar, & Sharp, 2015; Pandya-Jones, 2011; Perales & Bentley, 2009). Alternative splicing refers to the process by which exons of a gene may be included or excluded, producing numerous gene products (isoforms) and increasing gene product diversity and complexity (Black, 2003; Grabowski & Black, 2001). Both constitutive and alternative splicing occur co-transcriptionally, prior to the transcriptional termination and polyadenylation of pre-mRNAs (Pandya-Jones & Black, 2009). In addition to co-transcriptional splicing, there is also post-transcriptional splicing. It has been shown that the retention of individual introns in poly-adenylated pre-mRNAs serves as a mechanism for controlling gene expression; transcripts with retained introns will remain in the nucleus, preventing translation of the transcript, but following a cellular signal (e.g. osmotic or heat stress), the intron is excised and the protein is quickly translated (Boutz et al., 2015; Ninomiya, Kataoka, & Hagiwara, 2011). Here we closely studied *fgfr4*, a transcript that is both maternally deposited and zygotic transcribed and observe post-transcriptional splicing defects (in *fgfr4* as well as other targets) following the depletion of maternal and zygotic TAF15. Our RNAseq data, generated from post-ZGA embryos (stages 10 and 15) support a model whereby translation of maternally deposited *taf15* is required for the proper post-transcriptional splicing of nascent zygotic pre-mRNA transcripts during the time of zygotic genome activation. Indeed, in looking at early RNA seq datasets (Xenbase) we find no evidence that any of the maternal transcript (e.g. *fgfr4*) is incompletely spliced.

In the embryonic environment where zygotic genome activation occurs, and therefore active transcription is taking place, we propose that both maternally deposited TAF15, and TAF15 translated post-fertilization aids in splicing the new (zygotic) transcripts (e.g. *fgfr4, isl1*, and others; Supplemental Figures 2 and 4), resulting in unspliced transcripts at stage 10 after TAF15 depletion, but that zygotic TAF15 acts more classically and likely associates with the core promoter, using either its N-terminal low complexity domain or RNA-binding domains to bind the C-terminal domain of RNA pol II, and regulates transcription of specific targets (Figure 8). Upon zygotic genome activation (production of nascent transcripts), zygotic TAF15 may associate with the core promoter to regulate the expression of zygotically transcribed targets (e.g. *fgfr4*) (Figure 8). Our data also show that there is not always a discrepancy in maternal-zygotic TAF15 target regulation. In the case of *ventx2*.*1*, which is expressed zygotically (and not maternally deposited), we do not observe a splicing defect following TAF15 depletion, and find that both maternal and zygotic TAF15 depletion results in increased *ventx2*.*1* expression.

We now know that at least one member of the FET family of atypical RNA-binding proteins, TAF15, is required to regulate dorsoventral patterning in *Xenopus. Xenopus* embryos depleted of TAF15 have a phenotype similar to that which we observe when embryos are depleted of the BMP-antagonist *chordin*, or pairs of BMP antagonists (Reversade et al., 2005; Khokha et al., 2005). Both *chordin* and *taf15* depletion results in ventralized embryos with reduced head and eye structures as well as reduced dorsal and posterior fin and tail structures. Just as with TAF15 depletion, *chordin-*depleted embryos have a relatively normal cement gland and increased ventral tissue. Interestingly, according to our RNAseq data, TAF15-depleted embryos do not decrease *chordin (chrd)* expression, in fact we observe a two-fold increase at stage 15 (Supplemental Table 4 and 6). However, rather than regulate BMPs directly, we suggest a model where TAF15 is needed to repress *vent* genes and thereby disrupts the BMP/Vent positive feedback loop (Figure 6M) (Schuler-Metz et al., 2000).These data clearly illustrate the role of TAF15 in regulating dorsoventral patterning. We propose a model where TAF15 represses *ventx2*.*1* from the dorsal marginal zone of the gastrula and this repression continues through neurulation; we see TAF15 repressing *ventx2*.*1* from dorsal neural ectodermal tissue. Without knowing if the *ventx* genes are direct transcriptional targets of TAF15, we cannot conclude if the function of TAF15 is to repress *ventx* expression or to activate a repressor of *ventx* expression. However, the RNAseq results show that there is no intron retention in the *ventx* genes which supports the conclusion that the RNA splicing activity of TAF15 does not control the TAF15-dependent repression of *ventx2*.*1*.

Interestingly, the human Vent-like homeobox gene *VENTX*, a putative homolog of the *Xenopus ventx2* gene is aberrantly expressed in CD34+ cells of acute myeloid leukemia patients (Rawat et al., 2010). Furthermore, the leukemia-associated TAF15 fusion protein, TAF15-CIZ/NMP4, is found in acute myeloid leukemia (Alves et al., 2009). Although *ventx* has been lost in mouse, the function of *ventx* in repressing dorsal fates is well conserved between fish and frogs (Imai et al., 2001; Rawat et al., 2010). Given the relationship we have observed of increased *ventx* expression following TAF15 depletion, the coincidence of TAF15 dysfunction and increased *VENTX* in acute myeloid leukemia, and the fact that Ventx is required for proper mesenchyme and blood differentiation in *Xenopus*, it is possible that TAF15-dependent negative regulation of *ventx* is a conserved mechanism (Onichtchouk et al., 1998).

In summary, the data presented here show that a target gene (*fgfr4*) is regulated through two different molecular mechanisms (post-transcriptionally or transcriptionally) depending on the mechanism of TAF15 depletion (maternal + zygotic or zygotic-only). We have also demonstrated that while the effects of TAF15 on development are pleiotropic, consistent with our observations that embryos exhibit reduced head structures following TAF15 depletion, we have identified two specific pathways by which TAF15 regulates dorsoanterior brain development (*fgfr4* and *ventx2*.*1*). Furthermore, we have demonstrated that the FET family of atypical RNA-binding proteins do not act redundantly in regulating *Xenopus* development; TAF15 and Fus-depleted embryos exhibit very distinct phenotypes as well as mechanisms of gene expression regulation. Our findings of non-overlapping transcriptional effects by TAF15 and Fus in *Xenopus* supports the idea that the disease consequences resulting from chromosomal translocations involving the FET family of proteins are unlikely to be common transcriptional effects, but are more likely to result from the similar cellular aggregates of mutant proteins (Scekic-Zahirovic et al., 2016; Sharma et al., 2016; Svetoni et al., 2016).

## Supporting information

Combined supp figs

Supplemental Table 1. Differential Exon Usage, stage 10

Supplemental Table 2. Differential Exon Usage, stage 15

Supplemental Table 3. Stage-persistent Differential Exon Usage, Stage 10 and 15

Supplemental Table 4. Increased Gene Expression, stage 15

Supplemental Table 5. All changes in Gene Expression, stage 10

Supplemental Table 6. Increased and Decreased Gene Expression, stage 15

Supplemental table 7_intron retention characteristics

## Author Contributions

C.S.D. and RMH) designed and performed the experiments and wrote and edited the manuscript. D.D.S. assisted in bioinformatic analysis and laid the groundwork for studying FET proteins in *Xenopus*. C.R.T.E. and Y.X. performed embryo injections and imaging for revision experiments. R.M.H. conceived of the project and edited the manuscript.

## Acknowledgements

We would like to thank Marta Truchado Garcia and Edivinia Pangilinan of the Harland lab at the University of California, Berkeley for organizing reagents for revision experiments. We would also like to thank Helen Willsey of Matthew State’s lab for providing *Xenopus tropicalis* embryos for revision experiments. We would like to thank Jim Boonyaratanakornkit and Matthew Gray of Justin Taylor’s lab at the Fred Hutchinson Cancer Research Center for their time and Western blot reagents used for revision experiments.

## Competing Interested

The authors declare no competing financial interests.

## Notes

### Competing Interest Statement

The authors have declared no competing interest.

