## Supplementary material for "The atypical RNA-binding protein TAF15 regulates dorsoanterior neural development through diverse mechanisms in *Xenopus tropicalis*": Combined supp figs

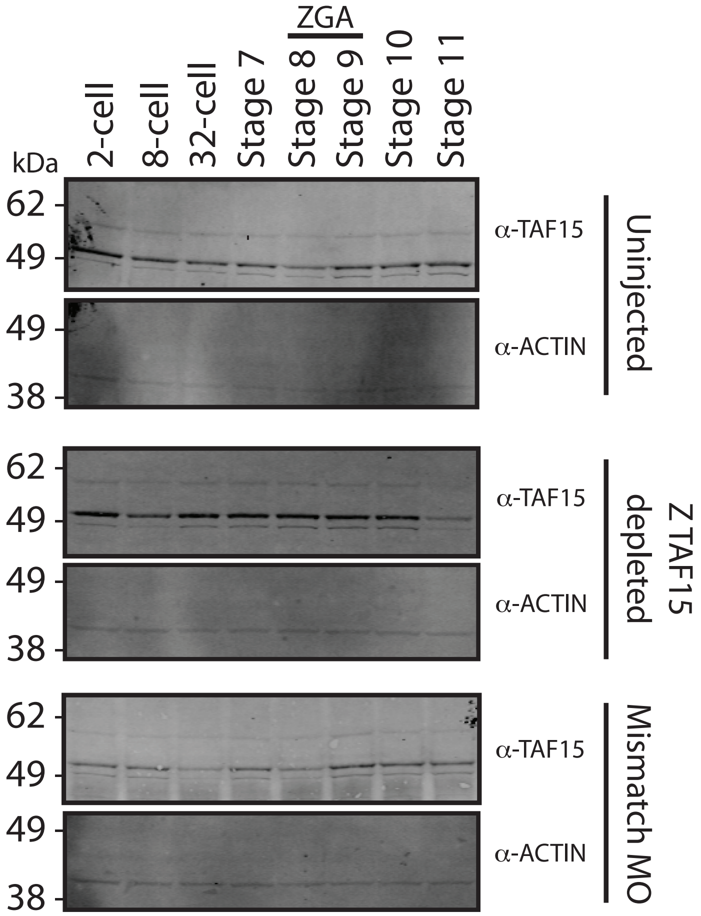
ZGA
2-cell
8-cell
32-cell
Stage 7
Stage 8
Stage 9
Stage 10
Stage 11
kDa
62
49
49
38
62
49
49
38
62
49
49
38
α-TAF15
α-ACTIN
Uninjected
α-TAF15
α-ACTIN
Z TAF15 depleted
α-TAF15
α-ACTIN
Mismatch MO

### Stage 10: top 10 named genes as sorted by adjusted P-value

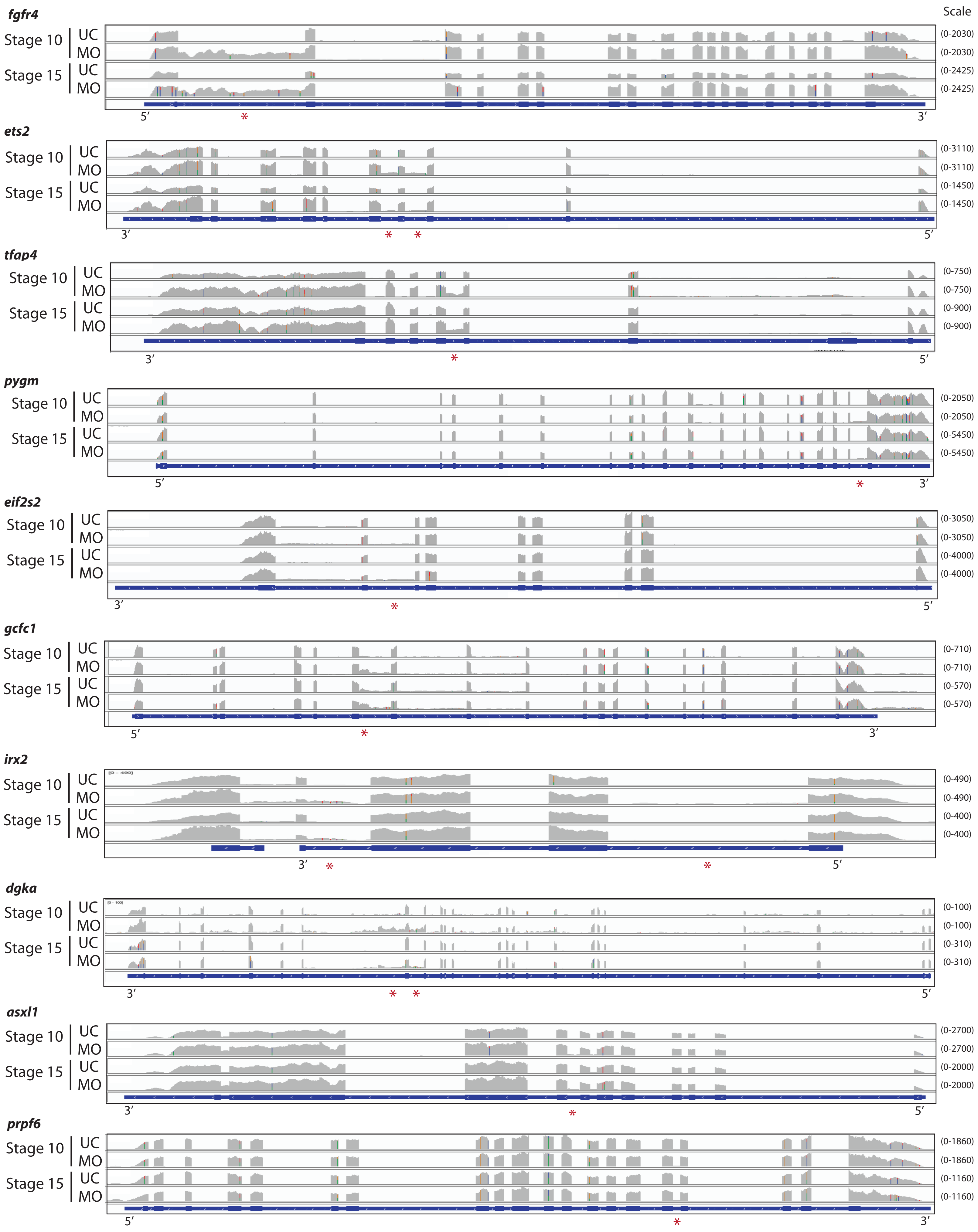


DeJong et al. Supplemental Fig 2

**Stage 15: top 10 named genes as sorted by adjusted P-value**

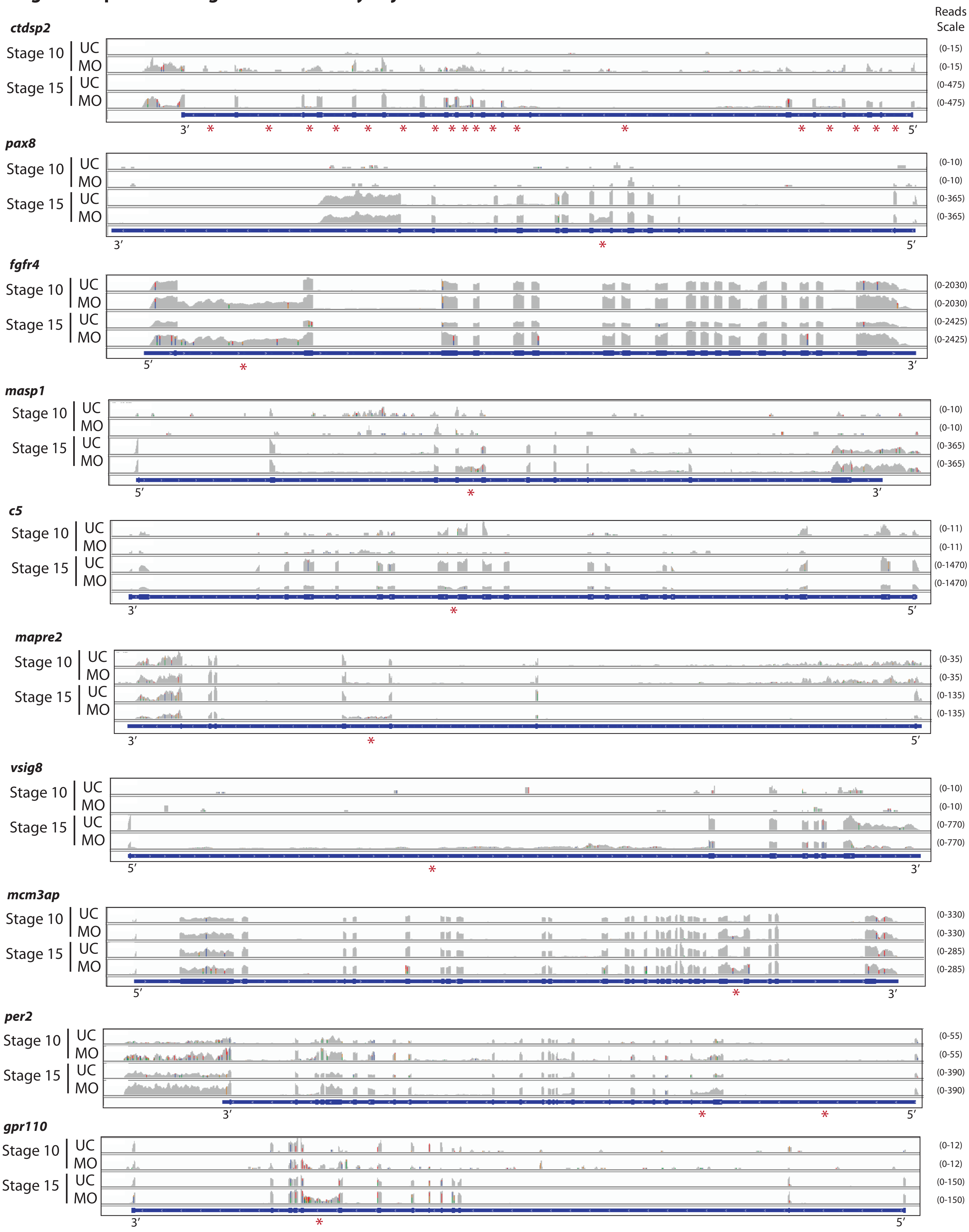


DeJong et al. Supplemental Fig 3

A

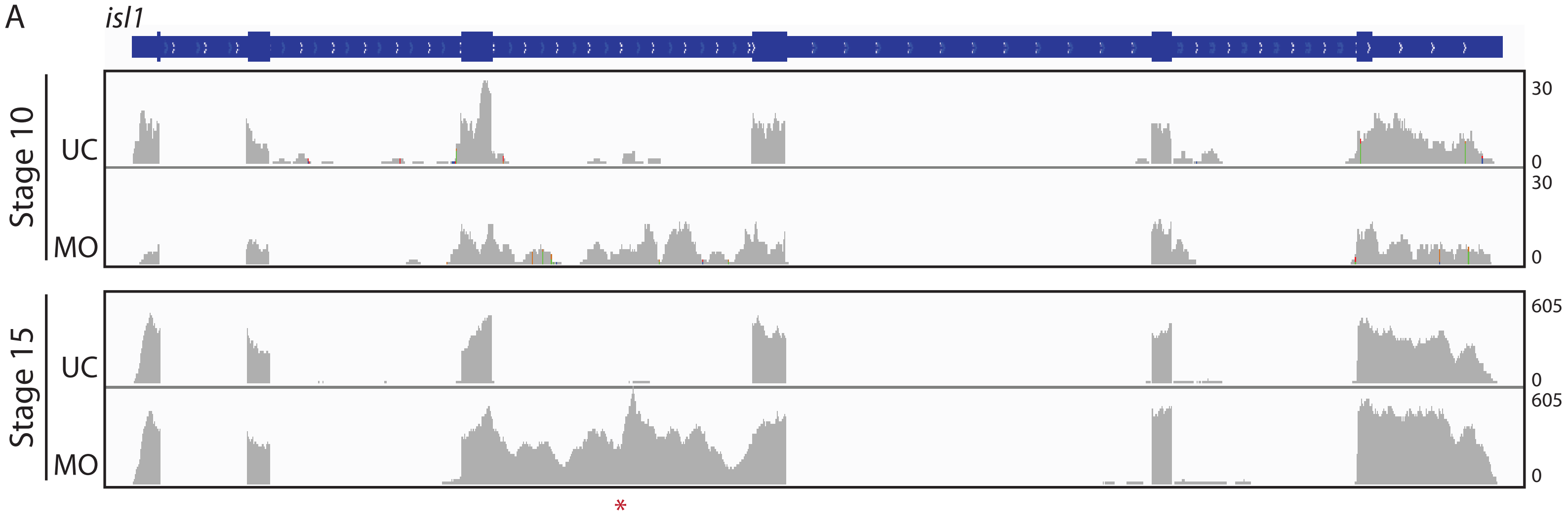

#### SUPPLEMENTAL FIGURE AND TABLE LEGENDS

**Supplemental Figure 1. TAF15 depletion by splice-blocking morpholino specifically targets zygotic *taf15*.** Western blot analysis of TAF15 in 2-cell (pre-zygotic genome activation) through stage 11 (post-zygotic genome activation) embryos from the following conditions: uninjected, splice-blocking morpholino (Z TAF15 depletion), and mismatch morpholino.

**Supplemental Figure 2. TAF15 depletion by translation-blocking morpholino leads to various types of intron retentions at stage 10.** Visualization of RNAseq reads with Integrative Genomics Viewer aligned with gene model in blue. Top 10 genes with differential exon usage (DEU) as sorted by their adjusted P value at stage 10. UC = uninjected control; MO = M+Z TAF15-depleting morpholino. * = intron with affected DEU according to DEXseq and does not indicate the number of DEU within an intron.

**Supplemental Figure 3. TAF15 depletion by translation-blocking morpholino leads to various types of intron retentions at stage 15.** Visualization of RNAseq reads with Integrative Genomics Viewer aligned with gene model in blue. Top 10 genes with differential exon usage (DEU) as sorted by their adjusted P value at stage 15. UC = uninjected control; MO = M+Z TAF15-depleting morpholino. * = intron with affected DEU according to DEXseq and does not indicate the number of DEU within an intron.

**Supplemental Figure 4. TAF15 depletion by translation-blocking morpholino leads to single intron retention in *ils1*.** (A) Visualization of *isl1* RNA-seq reads with Integrative Genomics Viewer aligned with gene model in blue. UC = uninjected control; MO = M+Z TAF15-depleting morpholino. * = intron with affected DEU according to DEXseq and does not indicate the number of DEU within an intron.

**Supplemental Table 1. Differential Exon Usage, stage 10.** DEXseq analysis of sequenced RNA from M+Z TAF15-depleted *X. tropicalis* embryos, stage 10; two-fold increase in expression cutoff; sorted by padj value.

**Supplemental Table 2. Differential Exon Usage, stage 15.** DEXseq analysis of sequenced RNA from M+Z TAF15-depleted *X. tropicalis* embryos, stage 15; two-fold increase in expression cutoff; sorted by padj value.

**Supplemental Table 3. Stage-persistent Differential Exon Usage, Stage 10 and 15.** Genes with conserved differential exon usage from stage 10 to 15. DEXseq analysis of sequenced RNA from M+Z TAF15-depleted *X. tropicalis* embryos; two-fold increase in expression cutoff; sorted by gene name.

**Supplemental Table 4. Increased Gene Expression, stage 15.** DESeq analysis of sequenced RNA from M+Z TAF15-depleted *X. tropicalis* embryos, stage 15; two-fold increase in expression cutoff; sorted by padj value.

**Supplemental Table 5. All changes in Gene Expression, stage 10.** DESeq analysis of sequenced RNA from M+Z TAF15-depleted *X. tropicalis* embryos, stage 10; sorted by padj value.

**Supplemental Table 6. Increased and Decreased Gene Expression, stage 15.** DESeq analysis of sequenced RNA from M+Z TAF15-depleted *X. tropicalis* embryos, stage 15; two-fold increase or decrease in expression cutoff; sorted by fold change.

**Supplemental Table 7. Differential Exon Usage Characteristics, stages 10 and 15.** Number and location of retained introns within the top 10 genes with differential exon usage (DEU) following DEXseq analysis, as sorted by adjusted P value.
