## Supplemental Table 3. Stage-persistent Differential Exon Usage, Stage 10 and 15 for "The atypical RNA-binding protein TAF15 regulates dorsoanterior neural development through diverse mechanisms in *Xenopus tropicalis*"

### DeJong et al. Supplemental Table 3. Stage-persistent Differential Exon Usage, Stage 10 and 15

| Gene | Condition UP | Retained intron as visualized by DEXseq gene modeling | Increased levels of intron detected as visualized by DEXseq gene modeling | Maternally deposited? | > 2 Fold increase? | # of genes |
|---|---|---|---|---|---|---|
| actr3 | MO | | x | yes | yes | |
| aldoa | MO | | x | yes | yes | |
| alms1 | MO | | x | yes | yes | |
| arhgap12 | MO | | x | yes | yes | |
| arrdc2 | MO | x | | yes | yes | |
| asxl1 | MO | | x | yes | yes | |
| bmp7.1 | MO | x | | yes | yes | |
| c11orf2 | MO | x | | yes | yes | |
| c16orf72 | MO | | x | yes | yes | |
| c1orf112 | MO | x | | yes | yes | |
| caprin1 | MO | | x | yes | yes | |
| cdc6 | MO | | x | yes | yes | |
| cdk12 | MO | | x | yes | yes | |
| cecr2 | MO | | x | yes | yes | |
| cep135 | MO | | x | yes | yes | |
| chmp5 | MO | x | | yes | yes | |
| clp1 | MO | x | | yes | yes | |
| ctdsp2 | MO | | x | yes | yes | |
| dennd2a | MO | x | | yes | yes | |
| dgka | MO | x | | yes | yes | |
| dido1 | MO | x | | yes | yes | |
| dnajc5g | MO | | x | yes | yes | |
| dus4l | MO | | x | yes | yes | |
| eed | MO | | x | yes | yes | |
| eif2s2 | MO | | x | yes | yes | |
| ets1 | MO | | x | yes | yes | |
| ets2 | MO | | x | yes | yes | |
| exosc2 | MO | | x | yes | yes | |
| extl3 | MO | | x | yes | yes | |
| farsa | MO | | x | yes | yes | |
| **fgfr4** | MO | x | | yes | yes | |
| fmnl3 | MO | | x | yes | yes | |
| gps2 | MO | x | | yes | yes | |
| gtf2h4 | MO | | x | yes | yes | |
| herpud1 | MO | x | | yes | yes | |
| hexdc | MO | x | | yes | yes | |
| hras | MO | | x | yes | yes | |
| ifrd2 | MO | | x | yes | yes | |
| ing4 | MO | x | | yes | yes | |
| ints6 | MO | | x | yes | yes | |
| irf6.2 | MO | | x | yes | yes | |
| jag1 | MO | | x | yes | yes | |
| kiaa0664 | MO | x | | yes | yes | |
| klf2 | MO | | x | yes | yes | |
| klhdc3 | MO | | x | yes | yes | |
| lemd3 | MO | | x | yes | yes | |
| lrig3 | MO | | x | yes | yes | |
| mcm3ap | MO | | x | yes | yes | |
| myd88 | MO | | x | yes | yes | |
| myo1e.2 | MO | | x | yes | yes | |
| neo1 | MO | x | | yes | yes | |
| nme2 | MO | | x | yes | yes | |
| npdc1.1 | MO | | x | yes | yes | |
| nxf1 | MO | | x | yes | yes | |
| pan3 | MO | x | | yes | yes | |
| parp1 | MO | | x | yes | yes | |
| pask | MO | | x | yes | yes | |
| paxip1 | MO | | x | yes | yes | |
| pdgfa | MO | x | | yes | yes | |
| per2 | MO | x | | yes | yes | |
| per3 | MO | | x | yes | yes | |
| pggt1b | MO | | x | yes | yes | |
| phf8 | MO | | x | yes | yes | |
| pip5k1a | MO | | x | yes | yes | |
| plekhg5 | MO | | x | yes | yes | |
| poldip3 | MO | | x | yes | yes | |
| polr2c | MO | | x | yes | yes | |
| prcp | MO | | x | yes | yes | |
| prpf39.1 | MO | | x | yes | yes | |
| prpf6 | MO | | x | yes | yes | |
| prr5l | MO | | x | yes | yes | |
| puf60 | MO | | x | yes | yes | |
| pus7 | MO | | x | yes | yes | |
| pygm | MO | | x | yes | yes | |
| rab15 | MO | x | | yes | yes | |
| rpf2:E002 | MO | | x | yes | yes | |
| rpl28 | MO | | x | yes | yes | |
| senp7 | MO | x | | yes | yes | |
| smpd1 | MO | | x | yes | yes | |
| spint2 | MO | | x | yes | yes | |
| srsf4 | MO | x | | yes | yes | |
| stam2 | MO | | x | yes | yes | |
| stoml2 | MO | | x | yes | yes | |
| tada1 | MO | x | | yes | yes | |
| tfap4 | MO | | x | yes | yes | |
| topbp1 | MO | | x | yes | yes | |
| trpc4ap | MO | | x | yes | yes | |
| ubap2 | MO | | x | yes | yes | |
| xnf7 | MO | | x | yes | yes | |
| xpo5 | MO | | x | yes | yes | |
| zdhhc5 | MO | x | | yes | yes | |
| znrf3 | MO | | x | yes | yes | 92 |

| | | | | | | |
|---|---|---|---|---|---|---|
| fam122b | MO | x | | no | yes | |
| frem3 | MO | | x | no | yes | |
| gpr110 | MO | x | | no | yes | |
| irx2 | MO | x | | no | yes | |
| isl1 | MO | x | | no | yes | |
| pdcd7 | MO | | x | no | yes | |
| phox2a | MO | | x | no | yes | |
| slc5a8 | MO | x | | no | yes | 8 |
| **Gene** | **Condition UP** | **Retained intron as visualized by DEXseq DEU model** | **Increased levels of intron detected as visualized by DEXseq DEU model** | **Maternally deposited?** | **> 2 Fold increase?** | **# of genes** |

DeJong et al. Supplemental Table 3. Stage-persistent Differential Exon Usage, Stage 10 and 15
