## Supplemental table 7_intron retention characteristics for "The atypical RNA-binding protein TAF15 regulates dorsoanterior neural development through diverse mechanisms in *Xenopus tropicalis*"

### DeJong et al. Supplemental Table 7. Differential Exon Usage Characteristics

#### Stage 10: top 10 named genes as sorted by adjusted P-value

| Gene | Number if retained introns | Which intron(s) retained? |
|---|---|---|
| fgfr4 | 1 | 1st |
| ets2 | 2 | 3rd and 4th |
| tfap4 | 1 | 3rd |
| pygm | 3 | Throughout 19th |
| eif2s2 | 1 | 7th |
| gcfc1 | 1 | 6th |
| irx2 | 4 | Throughout 1st and 3rd |
| dgka | 2 | 16th and 17th |
| asxl1 | 1 | 7th |
| prpf6 | 1 | 16th |

#### Stage 15: top 10 named genes as sorted by adjusted P-value

| Gene | Number if retained introns | Which intron(s) retained? |
|---|---|---|
| ctdsp2 | 22 | throughout whole transcript |
| pax8 | 1 | 5th |
| fgfr4 | 1 | 1st |
| masp1 | 1 | 4th |
| c5 | 1 | 12th |
| mapre2 | 1 | 3rd |
| vsig8 | 1 | 1st |
| mcm3ap | 2 | Throughout the 23rd |
| per2 | 2 | 1st and 2nd |
| gpr110 | 1 | 9th |
